# Variation in high-amplitude events across the human lifespan

**DOI:** 10.1101/2024.05.15.594378

**Authors:** Youngheun Jo, Jacob Tanner, Caio Seguin, Joshua Faskowitz, Richard F. Betzel

**Affiliations:** Department of Psychological and Brain Sciences, Indiana University, Bloomington, IN 47405; Cognitive Science Program, Indiana University, Bloomington, IN 47405; Luddy School of Informatics, Computing, and Engineering, Indiana University, Bloomington, IN 47405; Program in Neuroscience, Indiana University, Bloomington, IN 47405

## Abstract

Edge time series decompose functional connections into their fine-scale, framewise contributions. Previous studies have demonstrated that global high-amplitude “events” in edge time series can be clustered into distinct patterns. To date, however, it is unknown whether events and their patterns change or persist throughout the human lifespan. Here, we directly address this question by clustering event frames using the Nathan Kline Institute-Rockland sample that includes subjects with ages spanning the human lifespan. We find evidence of two main clusters that appear across subjects and age groups. We also find that these patterns of clusters systematically change in magnitude and frequency with age. Our results also demonstrate that such event clusters have distinct, heterogeneous relationships with structural connectivity-derived communication measures, which change with age. Finally, event clusters were found to outperform non-events in predicting phenotypes regarding human intelligence and achievement. Collectively, our findings fill several gaps in current knowledge about co-fluctuation patterns in edge time series and human aging, setting the stage for future investigation into the causal origins of changes in functional connectivity throughout the human lifespan.

## INTRODUCTION

Nervous systems are complex networks of anatomically connected neural elements–cells, populations, and areas linked by synapses, axonal projections, and myelinated white-matter tracts, respectively [1, 2]. The organization of these structural networks shapes brain-wide signaling patterns, inducing statistical dependencies between activity recorded from distant neural elements. Network science provides a mathematical framework for modeling both structural and functional connectivity (SC; FC), wherein neural elements are treated as nodes and their pairwise interactions as edges [3].

SC and FC undergo continuous and profound changes across the human lifespan [4–7]. Understanding this trajectory remains one of the central goals of neuroscience [8–10], promising insight into age-related changes in cognition and behavior [8, 11–13], neurodevelopmental disorders [14, 15], the progression of neurodegenerative disease [16], and neuropsychiatric conditions [17, 18]. More generally, tracking the normative trajectory of brain structure and function offers an invaluable reference for healthy brain function across the human lifespan [19].

Functional connectivity between brain regions is often summarized with measures of statistical dependence over time –e.g. correlation, coherence, mutual information. While static functional connectivity provides a time-invariant summary of statistical relationships between brain regions, connectivity is thought to fluctuate across time [20, 21]. Typically, sliding-window [22] and kernel-based approaches [23] are used to obtain time-varying estimates of FC. However, both approaches create an aggregate measure that spans multiple time points which may result in artificially smooth functional connectivity at fine-scale temporal resolution.

Recently, we showed that static FC could be decomposed into its framewise contributions, yielding timevarying estimates of coupling weights for each pair of nodes – so-called “edge time series.” [24, 25]. This approach builds upon existing frameworks to track the temporal dynamics in the brain’s functional connectivity [26–30]. Using edge time series, we identified brainwide “events”–intermittent and brief moments of global high-amplitude co-fluctuations [24, 25]. These events contribute disproportionately to static FC [24], can be predicted from static functional connectivity [31], may provide biomarkers for disorders (e.g. early mild cognitive impairment) [32], carry subject-specific information and enhance brain-behavior associations [33–36]. Events can also be partitioned into recurring “states” [33], whose topography and relative frequency may be involved with fluctuations of endogenous hormones [37].

While various studies have applied the “edge-centric” approach to human and even non-human imaging data [38, 39], investigations into how such high amplitude cofluctuations differ across the human lifespan has not yet been studied. For instance, it is unclear whether topographically similar events manifest in older and younger individuals or whether event topography varies with age. Furthermore, the relative frequency of events as a function of age is unclear. Additionally, little is known about the link between the underlying SC and events [38–41].

Here, we investigate events throughout the human lifespan using resting-state fMRI data from 537 subjects, spanning ages 6 to 75, from the Nathan Kline Institute - Rockland enhanced sample [42]. Using events detected in edge time series from subjects across age groups, we aimed to address the following questions. How do events and their patterns differ across ages throughout the human lifespan? Are event co-fluctuation patterns related to structural connectivity across development and aging? How are event co-fluctuation patterns organized and are they useful for making phenotypic predictions? Progress towards answering these questions may help clarify the possible drivers in normative changes in functional connectivity across the human lifespan.

## RESULTS

### Events can be clustered into distinct patterns

In this paper, we aimed to uncover patterns of events from data acquired across the human lifespan. Specifically, we used the resting-state fMRI data from the Nathan Kline Institute-Rockland sample with 537 subjects spanning ages 6 to 75 years [42] after excluding subjects with high-motion frames or missing imaging or metadata (detailed exclusion criteria in Methods). In all analyses, we used the Schaefer-Yeo parcellation [43] with *N* = 400 parcels to define the nodes in both functional and structural networks.

One of the challenges of working with the NKI dataset is its uneven distribution of participants’ ages. To reduce related biases, we aimed to create subsamples of the dataset in which ages are (approximately) uniform. To do this, we assigned each participant to one of seven age groups and sampled an equal number of subjects from each group (20 subjects per sampling procedure with replacement; Fig.1*a-b*). We then transformed the regional fMRI BOLD time series into edge time series (ETS) by calculating the element-wise product between all pairs of z-scored time series (Fig. 1*c*). An event detection algorithm was applied to the edge time series as described in previous literature [44, 45]. In brief, these events are detected by first identifying frames that surpass the co-fluctuation magnitudes of a null model created by circularly shifting each region’s activity time series (Fig. 1*d*). This null model preserves the mean and variance of regions’ activity while destroying correlation structures. We then performed a k-means clustering on this samples’ events (Fig. 1*f*). We repeat this sampling and clustering process (Fig. 1*b-f*) for 100 iterations at each K. Lastly, since the k-means algorithm produces slightly different cluster results in each iteration, all reported cluster results were aligned to that of the most similar centroid across all iterations.

**FIG. 1.**
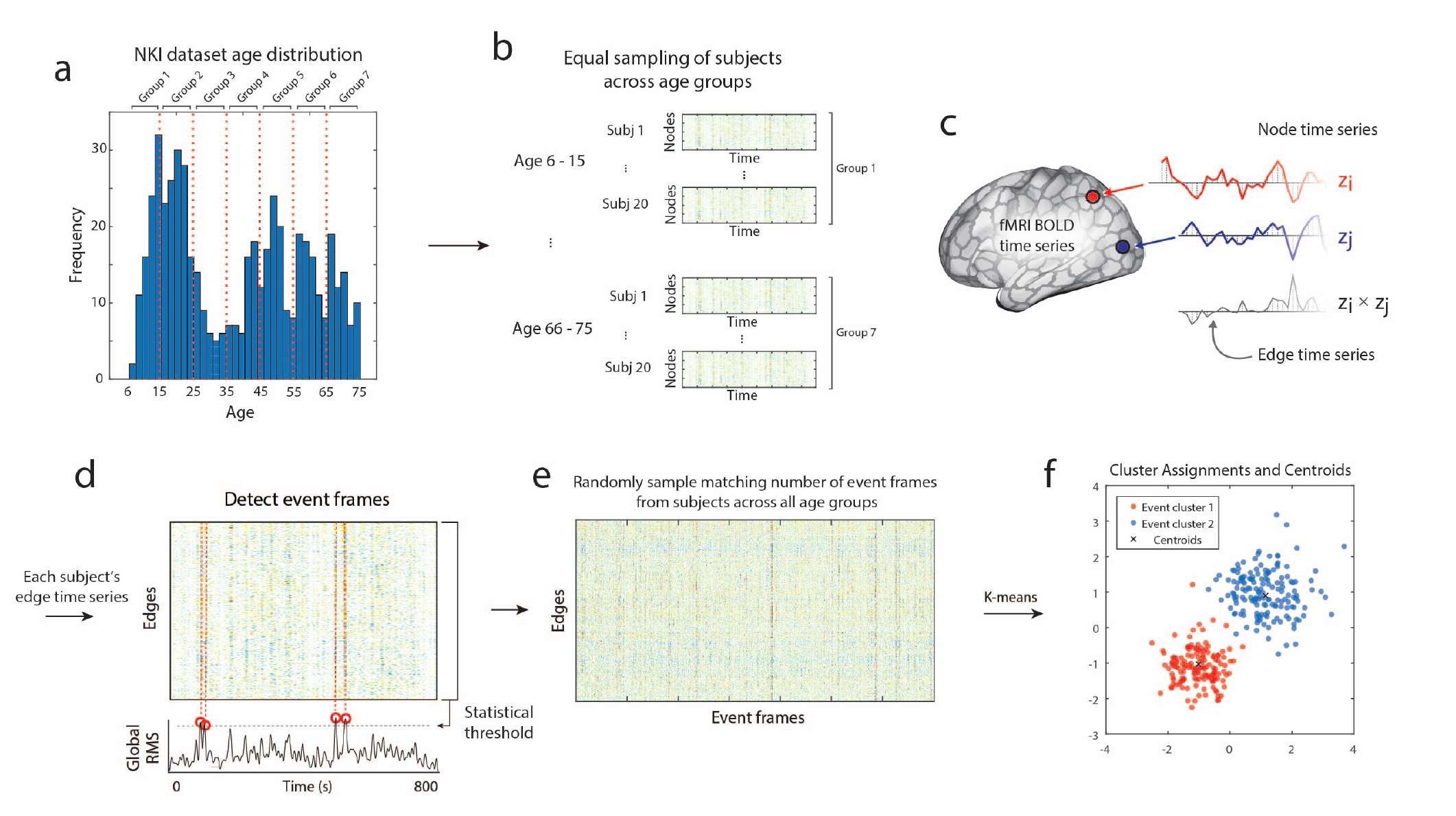
Schematic illustration of creating event clusters created with an equal number of events across age groups. (*a*) The age distribution of the NKI dataset (red lines: age bin boarders). (*b*) Matched number of subjects (*n* = 20) sampled per age group (seven age bins). (*c*) Edge time series calculated as the moment-to-moment multiplication of node time series. (*d*) Detection of “event” frames in edge time series by selecting frames above a statistical threshold. (*e*) Event frames of all subjects across all age groups on which we applied (*f*) k-means clustering to assign events to clusters and the cluster centroids.

With the events detected across all subjects and age groups (width of age bin = 10 years), we identified clusters at *K* = 2 − 10 using k-means clustering. Here, we used two distance metrics, the bivariate product-moment correlation coefficient and Lin’s concordance, to compare pairwise similarities for detecting clusters of events. The standard product-moment correlation rescales patterns (z-score) before computing similarity, whereas Lin’s concordance allows vectors to be distinguished from one another if their magnitudes vary [46]. We note that maximum similarity across clustering iterations were found at *K* = 2 for both distance metrics when aiming to minimize a cost function using the Hungarian algorithm (Fig. 2*a*). The resulting event clusters at *K* = 2 are shown in Fig. 2*b-c* (mean event cluster patterns across all subjects, from all age groups, and all iterations post-alignment). Additionally, similar clusters of events were also found using a spectral clustering algorithm (Fig. S9).

**FIG. 2.**
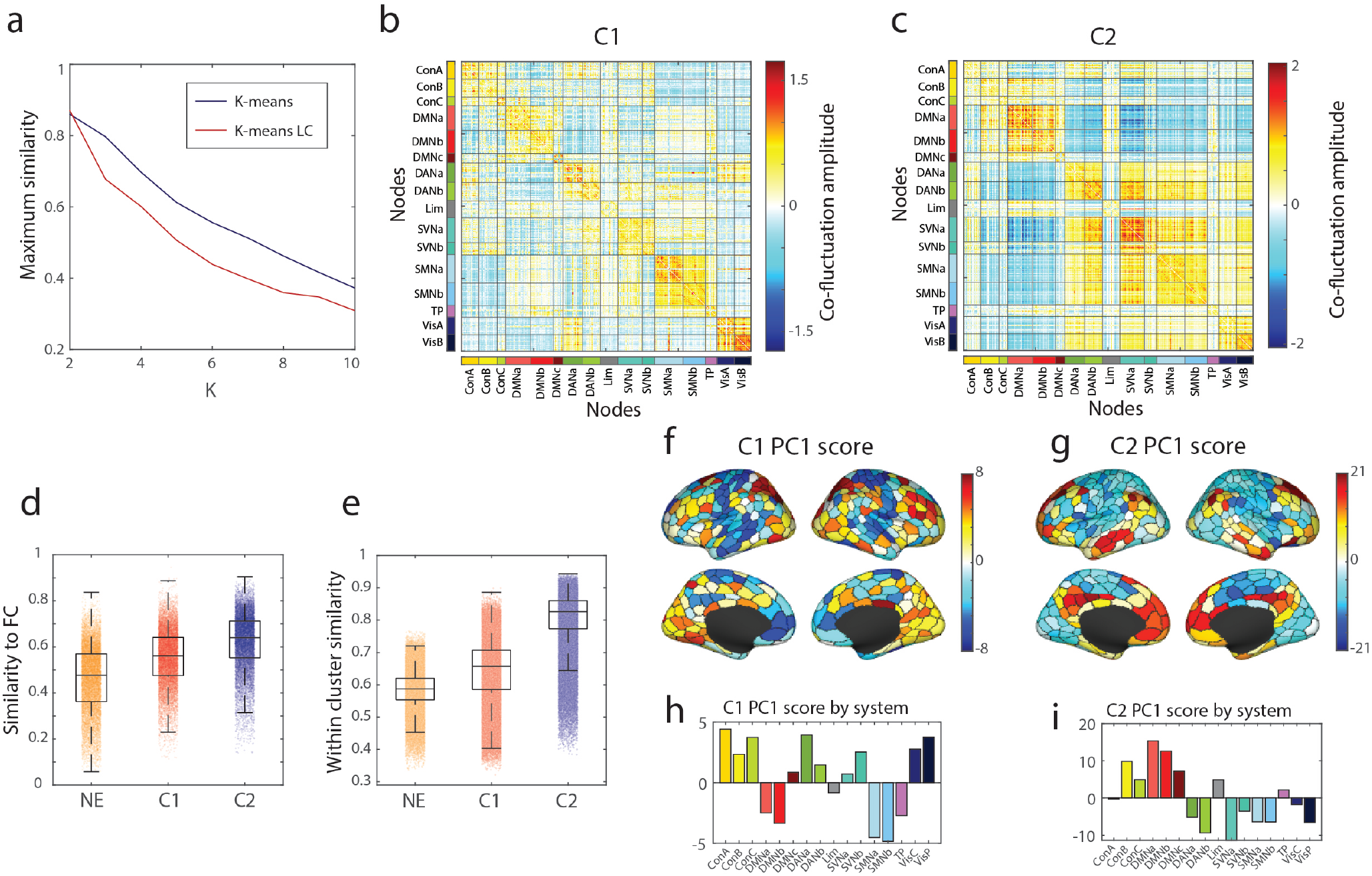
Event clusters 1 and 2 have distinct patterns and characteristics. (*a*) The maximum similarity across 100 runs of k-means clustering using Pearson correlation and k-means clustering using Lin’s concordance as a measure of similarity (*K* = 2 − 10). (*b*) Event co-fluctuation pattern cluster 1 at *K* = 2 averaged across all age groups and all runs. (*c*) Event co-fluctuation pattern of cluster 2 at *K* = 2 averaged across all age groups and all runs. (*d*) Similarity of C1 and C2 to static FC. (*e*) Scores of the first principal component of cluster 1. (*f*) Scores of the first principal component of cluster 2.

After detecting clusters, we described the event co-fluctuation patterns based on their similarity to FC, within-cluster homogeneity, and principal components. To do so, we first calculated each subject’s average matrix of cluster 1, cluster 2, and non-event (NE) frames to compare to the subject’s static FC matrix. Compared to cluster 1, cluster 2 was more strongly correlated with static FC (paired sample t-test; *p* < 10^−15^; Fig. 2*d*). Nevertheless, cluster 1 was more strongly correlated with static FC than NE frames (paired sample t-test; *p* < 10^−15^; Fig. 2*d*). Also, using the subject-level averages of cluster 1, cluster 2, and NE patterns, we found that the within-cluster similarity was greatest for cluster 2 compared to cluster 1 (*p* < 10^−15^) or non-events (*p* < 10^−15^; Fig. 2*e*).

Next, in order to investigate system-level differences in event co-fluctuation patterns, we applied principal component analysis to the subject-level averages of event co-fluctuation patterns and non-events. Notably, the first principal component (PC) of cluster 1 explained 41.7% of variance (Fig. 2*f*) compared to 88.0% of variance explained using PC1 in cluster 2 (Fig. 2*g*). When compared against canonical brain systems [43], the PC1 scores of cluster 1 loaded positively onto the control, dorsal attention, salience/ventral attention, and visual networks (Fig. 2*h*). Negative loadings in cluster 1 were found in the default mode, limbic, somatomotor, and temporoparietal networks. In cluster 2 (Fig. 2*i*), the scores in PC1 mainly revealed positive loadings in higher-order, heteromodal networks - control, default mode, limbic, and temporoparietal networks. In contrast, negative loadings were found in the unimodal networks in the dorsal attention, salience ventral attention, somatomotor, and visual networks.

In sum, cluster 2 better represented static FC, with greater within-cluster similarity, with significant alignment to the S-A axis, and was found to load positively on higher-order networks and negatively on unimodal networks. Cluster 1 was found to be less similar to static FC, with reduced within-cluster similarity, and was not aligned with the S-A axis, with mixed loadings on higher-order and unimodal networks. Together, these results indicate that events can be grouped into 2 clusters with distinct relationships to the brain’s functional architecture.

### Events patterns change with age

In the previous section we used a data-driven approach to identify two event co-fluctuation patterns that appear consistently across the lifespan. It remains unclear, however, whether these two patterns persist unchanged across age groups. That is, do these events occur at different frequencies in younger brains compared to older? Do changes in event co-fluctuation patterns occur heterogeneously across functional systems? In this section, we aim to address these questions.

First, we created the average event co-fluctuation pattern for each age group for both cluster 1 and cluster 2 and created a difference matrix between an age group of interest minus that of the youngest age group (Fig. 3*a*). Next, we used the first principal component of the difference matrix between the youngest and oldest age groups for cluster 1 (Fig. 3*b*) and cluster 2 (Fig. 3*c*) to visualize the difference in event co-fluctuation patterns across age groups on the nodal level.

**FIG. 3.**
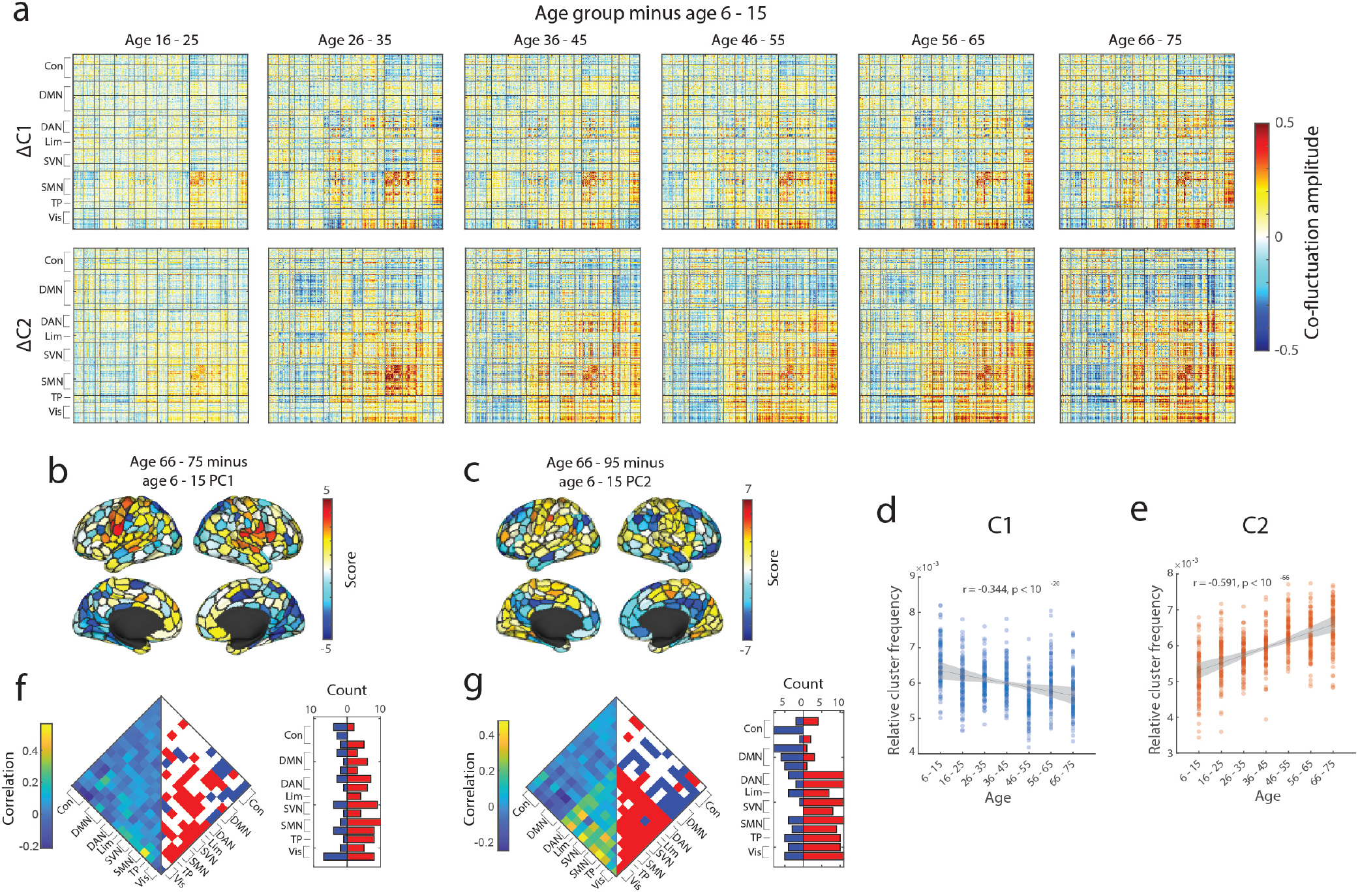
Differences in event clusters with age. (*a*) Cluster patterns were averaged for each cluster across all events and runs, and subtracted the youngest age group from the rest of the age groups to compare the average difference in event co-fluctuation patterns (top row: cluster 1; bottom row: cluster 2). (*b*) Scores of first principal component between the youngest and oldest age groups for cluster 1. (*c*) Scores of the first principal component between the youngest and oldest age groups for cluster 2. (*d*) Relative frequency of cluster 1 out of 10 randomly sampled event frames for each subject across age groups. (*e*) Relative frequency of cluster 2 out of 10 randomly sampled event frames for each subject across age groups. (*f*) System edges in cluster 1 and their correlation with age (left: correlation coefficients; right: significant systems below *p*_*adjusted*_ = 10^−3^). (*g*) System edges in cluster 2 and their correlation with age (left: correlation coefficients; right: significant systems below *p*_*adjusted*_ = 0.0061).

Next, we tested whether the frequency of event co-fluctuation patterns changes with age. Due to the variability in total scan duration and number of events across subjects, we randomly selected 10 event frames per subject prior to clustering the events. To calculate the relative frequency of each event co-fluctuation pattern, the total number of each event co-fluctuation pattern was divided by the total number of frames used after removing high-motion frames. We found that the relative frequency cluster 1 to significantly become less frequent with age (Fig. 3*d* ; *r* = −0.34; *p* < 10^−15^) whereas the occurrences of cluster 2 significantly increased with age (Fig. 3*e*; *r* = 0.59; *p* < 10^−15^).

We further investigated whether co-fluctuation patterns differ with age in event co-fluctuation patterns at the system level. To do so, we first created an average cluster 1 and cluster 2 matrix for each age group - averaging across subjects and iterations after aligning events to the cluster centroids. The edges that fall within or between particular systems were averaged, yielding a system-by-system matrix, and the elements of this matrix correlated with age. We then compared the observed correlations to that of an age-randomized null distribution which consisted of event clusters by system correlated with randomly re-assigned age groups (5000 iterations). Compared to the age-randomized null, we found system pairs whose mean co-fluctuation amplitude was significantly more correlated with age in cluster 1 and cluster 2 (Fig. 3*f-g*, FDR-adjusted p-values; *q* = 0.01; cluster 1 *p*_*adjusted*_ = 0.0037; cluster 2 *p*_*adjusted*_ = 0.0061). We found more significant system-level changes with age in cluster 2 than in cluster 1. Specifically, cluster 2 showed a bipartite pattern in correlations with age that were largely negative in the control and default mode networks versus positive age correlations in the dorsal attention, limbic, salience ventral attention, somatomotor, temporoparietal, and visual networks (Fig. 3*g*).

Lastly, we tested whether event co-fluctuation patterns and their properties generalize across different numbers of age bins of 5 and 10 years (Fig. S1). When using different numbers of age bins, we also found two clusters, which also showed clusters which were highly correlated with those of 7 age bins (7 age bins versus 5 age bins cluster 1 : *mean* = 0.82*±*0.12; cluster 2: *mean* = 0.92*±*0.076; 7 age bins versus 10 age bins cluster 1: mean = 0.86*±*0.11; cluster 2: mean = 0.94 *±* 0.065; Fig. S1*c, g*). We also found the within-cluster similarity in cluster 2 to be greater than cluster 1 in both 5 and 10 age bins (5 age bins *p* < 10^−15^; 10 age bins; *p* < 10^−15^) as in the results with 7 age bins. The pattern of age-related changes in relative frequencies of each cluster also matched the results of 7 age bins (Fig. S2; cluster 1: *r* = −0.37; *p* < 10^−15^; cluster 2: *r* = 0.62; *p* < 10^−15^).

In summary, we found that event co-fluctuation patterns change with age in both their frequency and their system-level organization. Cluster 1 became less frequent with age, which revealed a heterogeneous system-level change with age. Cluster 2 frequencies increased with age, with decreased co-fluctuation within higher-order networks and increased co-fluctuation patterns in unimodal networks with age. Together, these results suggest that changes in the organization of functional connectivity across the human lifespan may be highlighted and further dissected when focusing on the changes in patterns of events - effects of which may be obscured when averaging across the entire time series data.

### Local SC-based communication measure-event cluster coupling changes with age

Understanding the relationship between brain structure and function is a central goal in neuroscience. Typical SC-FC studies allows one to investigate the association between structural and functional connection weights, which does not allow the researcher to investigate polysynaptic interactions known to shape brain function. Network communication models are a framework to quantify the structural capacity for interregional communication in the connectome, which takes into account not only direct connections but also putative polysynaptic paths [47]. These models have been used to investigate SC-FC coupling in static FC [48–51], across the human lifespan [40, 52], to investigate propagation of electrical stimulation [53]. Here, for the first time, we use communication models to understand how the structural basis of event co-fluctuation pattern organization changes with age.

In brief, these communication models can be largely organized based on whether the measure’s policy aims to explain communication as a “centralized” or “decentralized” process. For instance, the measure of shortest paths is a “centralized” communication measure in that the signaling process requires the knowledge of the entire network’s topology. On the other hand, diffusive communication policies such as random walks [54] are “decentralized” policies since the policy is dependent on the information available at each node rather than the topology of the complete network.

Here, we aim to answer the following questions by applying communication models to the lifespan data. If we generate communication policies based on structural connectivity, can we explain variation in event co-fluctuation patterns throughout the human lifespan? Does the relationship between each event co-fluctuation pattern and structural connectivity-derived communication measures change with age? To address these questions, we used 9 different communication measures (and their weighted variants totaling 34 policies) that embody various policies of network communication and Euclidean distance to link structural connectivity with event co-fluctuation patterns [40].

Here, we used each subject’s SC matrix to create 34 matrices embodying different communication policies (Fig. 4*a-b*). We then used the average event co-fluctuation pattern for cluster 1 and cluster 2 for each age group (Fig. 4*c*) to calculate the variance explained by the communication measure for each node in the event co-fluctuation patterns (Fig. 4*d*). We found the overall SC-to-cluster relationship to vary across nodes when averaging the explained variance across communication measures (Fig. 4*e*). To identify which functional system is best explained by the communication models, we calculated the maximum variance explained per node (Fig. 4*f-g*) and summarized the variance explained by functional system (Fig. 4*h-i*). In cluster 1, the maximum variance explained across various communication measures was mainly in the visual, dorsal attention, and somatomotor networks (Fig. 4*f, h*). The maximum variance explained in cluster 2 was in the default A, default B, control B, control C, limbic, somatomotor A, and visual systems (Fig. 4*g, i*). Highlighted systems in Fig. 4*h, i* had significantly greater explained variance than that of the nodal spin test permutation of 5000 iterations (*p*_*adjusted*_ < 0.0002).

**FIG. 4.**
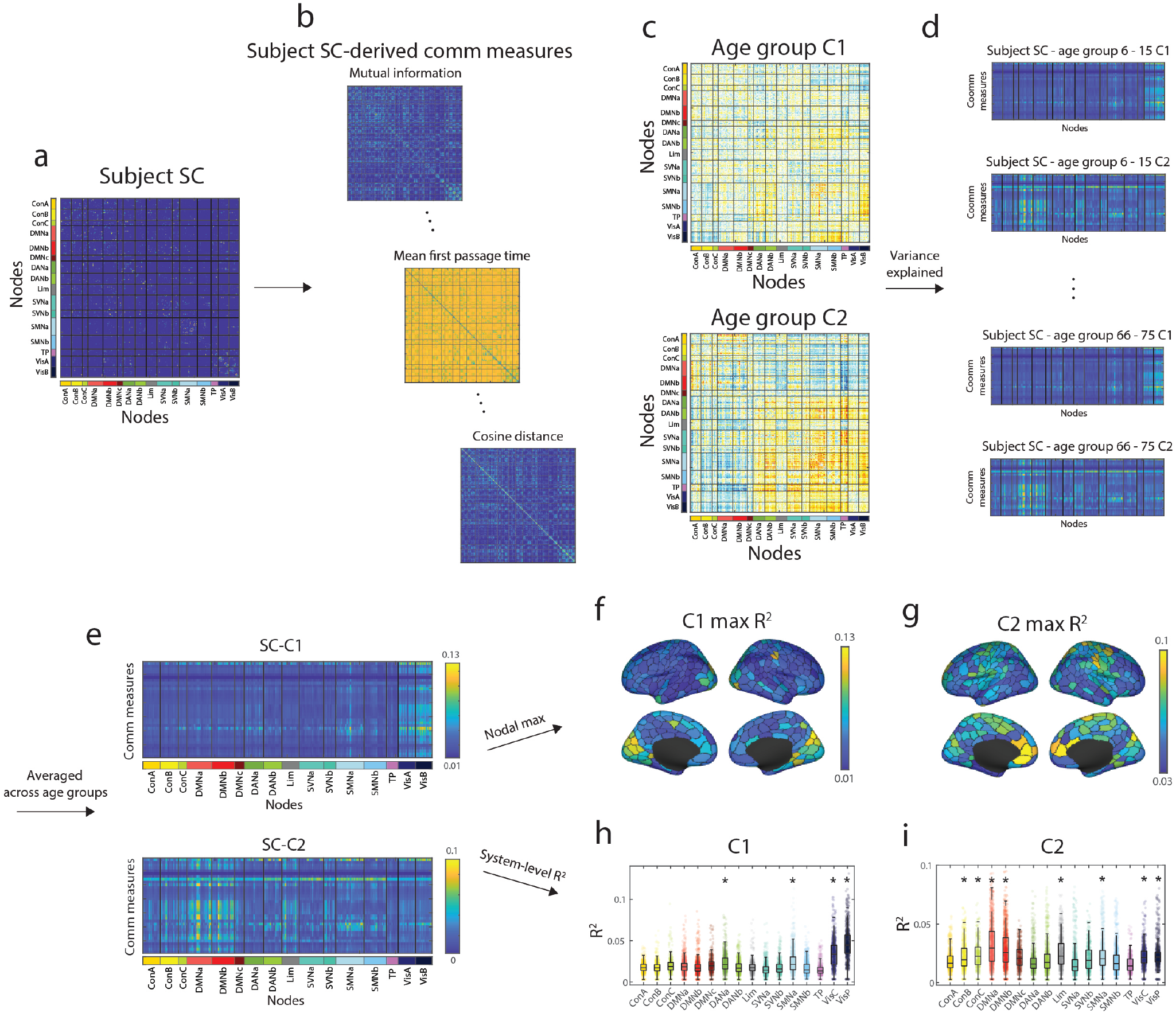
Schematic illustration of calculating the explained variance between event clusters and structural connectivity-derived communication measures. We take a subject’s structural connectivity (SC) matrix (*a*) to create 34 different measures that incorporate various communication policies. Next, we create the average cluster 1 and cluster 2 events for each age group (*b*). Matrices from steps (*b-c*) were used to calculate the variance explained for each node in cluster 1 and cluster 2 by each communication measure (*d*). We calculated the mean event co-fluctuation pattern-to-SC-derived measure relationship matrix for cluster 1 and cluster 2 across all age groups (*e*). This can be used to find the maximum variance explained per node for cluster 1 (*f*) and (*g*) or plot the maximum variance explained by system (*h-i*).

After investigating the relationship of event co-fluctuation patterns with SC-derived communication measures, we then asked whether these relationships change with age. To answer this question, we calculated the variance explained in event co-fluctuation patterns by communication measures for each age group. Next, we narrowed the scope of investigation by identifying communication measures that best explained the variance of cluster centroids across nodes for cluster 1 and cluster 2 (Fig. 5*a-b*). We also showed the system-level patterns in subject-level SC-derived communication measures and how their event co-fluctuation pattern relationships vary across age groups (Fig. 5*c*). These results were also found in distance-dependent, representative SC matrices for each age group and event co-fluctuation patterns (Fig. S10, [55]).

**FIG. 5.**
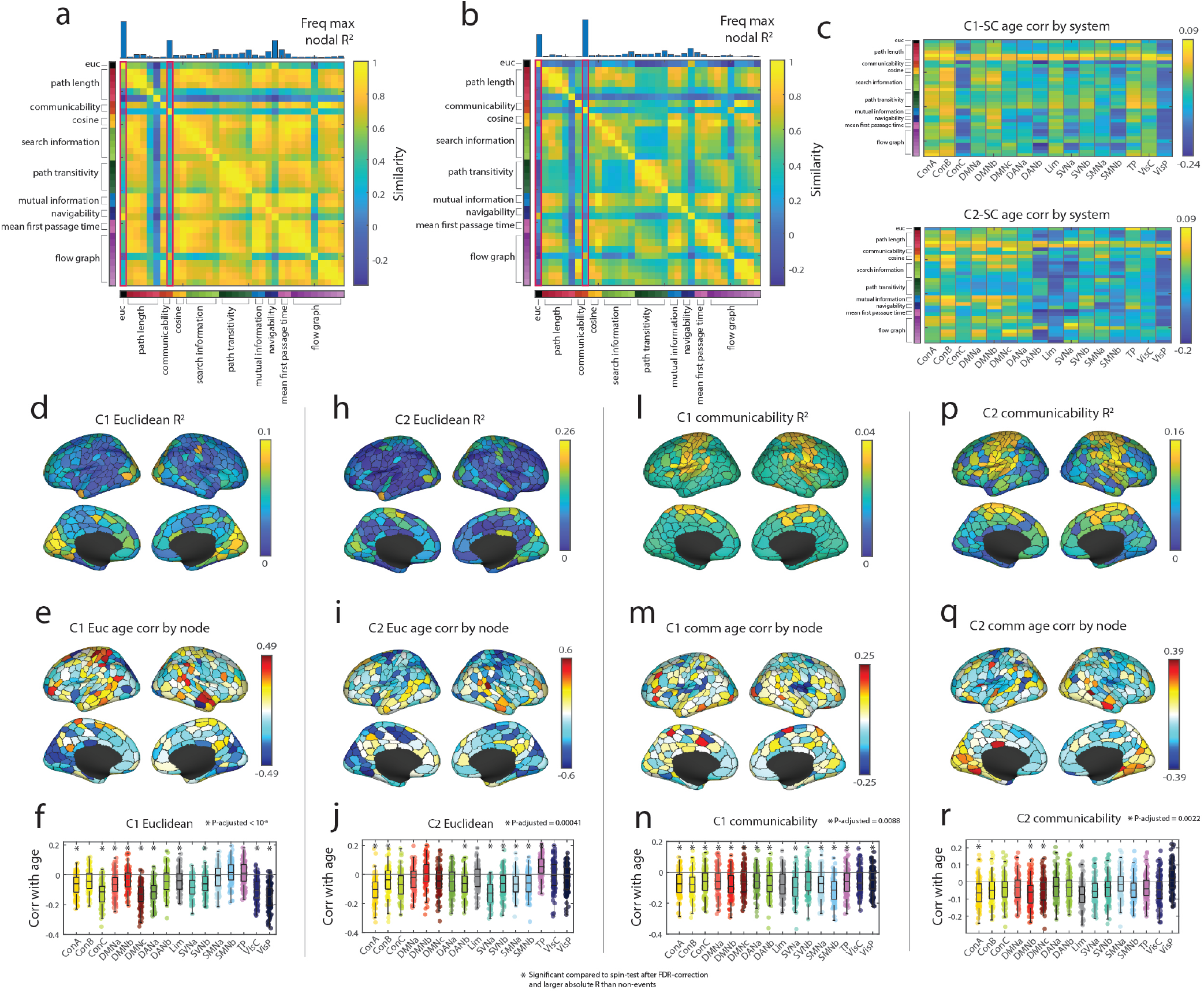
SC-event relationships and age. (*a*) Similarity of communication measures for cluster 1 events. Histogram on top represents the count of each communication measure having the maximum nodal explained variance. (*b*) Similarity of communication measures for cluster 2 events. Histogram on top represents the count of each communication measure having the maximum nodal explained variance. The two measures exhibiting greatest *R*^2^ were Euclidean distance and communicability, highlighted in magenta boxes. (*c*) System-level correlations of cluster 1 (top) and cluster 2 (bottom) events with various SC-derived communication measures with age. Panels *d, e* and *f* show variance in the cluster 1 centroid explained by Euclidean distance, its correlation with age, and the breakdown of these correlations by brain system. The remaining panels show similar information for cluster 2 and Euclidean distance, and clusters 1 and 2 with communicability.

In both cluster 1 and cluster 2, Euclidean distance was found as one of the most frequently maximal measures of explained variance. The explained variance of both cluster 1 and cluster 2 events using Euclidean distance was highlighted in the visual networks (Fig. 5*d, h*). When taking the explained variance of cluster 1 events by Euclidean distance and correlated them across age groups, we only found negative significant correlations with age in the control A, control C, default mode, dorsal attention A, salience ventral attention, and visual networks (Fig. 5*e-f*). Here, we considered the system-level age correlation significant if the correlation was both significant compared to a spin-test null model and the magnitude of age correlation greater than that of non-event frames (Fig. 5*f* ; *p*_*a*_*djusted* < 10^−8^). When correlating the explained variance of cluster 2 events and Euclidean distance with age, we found significantly positive correlations only in the temporoparietal network, and significant negative correlations in the control A and B, dorsal attention B, salience ventral attention, and somatomotor networks (Fig. 5*i-j* ; *p*_*adjusted*_ = 10^−4^).

Another frequent measure of maximum explained variance in cluster 1 and cluster 2 was communicability - which gives more weight to shorter walks in a network that accounts for various paths between two nodes [56]. With communicability, in both clusters 1 and 2, the explained variance was mainly enriched in the somatomotor networks (Fig. 5*l, p*). The explained variance of cluster 1 events and communicability was found to have significant negative correlations across most systems except the limbic, salience ventral attention B, and central visual networks (Fig. 5*m, n*; *p*_*a*_*djusted* = 0.0088). When correlating the explained variance of cluster 2 events and communicability with age, we found significantly negative correlations in the control A, default mode B, default mode C, and limbic networks (Fig. 5*q, r*).

Combined, we find the results in event patterns to align with previous research that Euclidean distance and communicability explain structure-function relations in static FC [40]. Also, most age correlations between event co-fluctuation patterns on the system-level with SC-derived communication measures were negatively correlated. In other words, previous work has showed that structure-static FC relations globally decrease with age but variable in canonical brain networks [40], we were also able to observe a similar phenomena when using patterns of events with age. However, the extent of different measures’ and their system-level explained variances were heterogeneous across the event co-fluctuation patterns.

### Modular event co-fluctuation patterns

In this section, we investigated how the event co-fluctuation patterns are organized - i.e. do events depict modular structure as static FC? If so, how are their modules organized and do they decrease with age as reported in previous studies using static FC? In this section, we aimed to address these questions by using modularity maximization with generalized Louvain heuristics. We calculated modularity (*Q*) in the event clusters, non-event frames, and static functional connectivity.

Here, we show that cluster 1 and cluster 2 events are significantly more modular than non-events across age groups, and that the modularity of static FC falls in between events and non-events (Fig. 6*a*). Our results also show that modularity of both event co-fluctuation patterns decrease with age and that the modularity of static FC is driven by the modularity of event frames. These results also align with previous literature demonstrating how the modularity of static FC decreases with age. Next, we weighed each event co-fluctuation pattern’s modularity with their frequencies in each age group (Fig. 3*d-e*). This allows a more realistic estimate of the combined impact of changes in event co-fluctuation pattern modularity and how they overall change with age. Here, we found significant decreases in modularity with age when weighing events with their frequencies (Fig. 6*b*; *r* = −0.48; *p* < 10^11^).

**FIG. 6.**
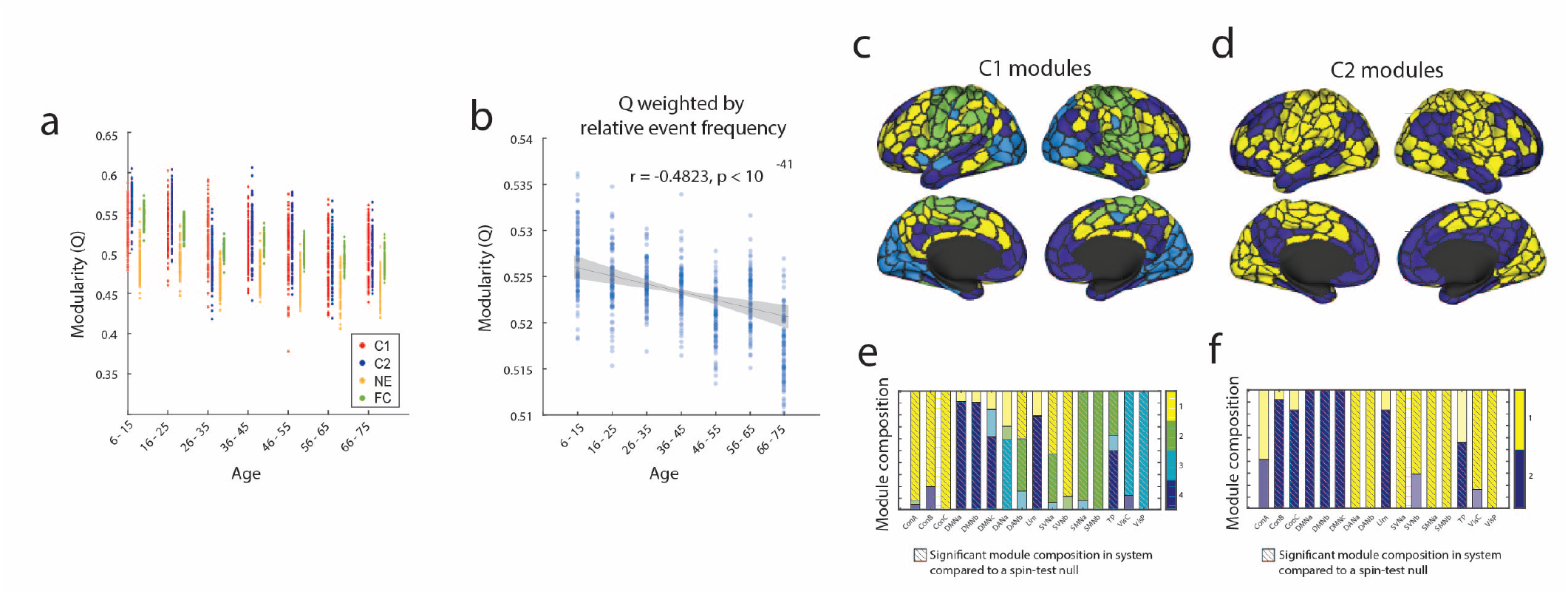
Event modules. (*a*) Modularity of event co-fluctuation patterns, non-events, and static FC across age groups. (*b*) Modularity of event co-fluctuation patterns after weighing each pattern with its frequency in each age group. Modules of (*c*) cluster 1 and (*d*) cluster 2 and their system-level composition (*e-f*). Modules highlighted in magenta boxes are modules that comprise a significantly greater proportion of the functional system than a spin-test null result of 5000 iterations.

Next, we asked whether there were differences in the modular organizations between the event co-fluctuation patterns. When using modularity maximization in cluster 1 events across all age groups and subjects, we detected four modules (Fig. 6*c, e*). When comparing the module compositions to that of a nodal spin-test null of 5000 iterations, we found control, dorsal attention B, salience ventral attention networks to be represented above change in module 1, the limbic, somatomotor, temporoparietal networks in module 2, the visual and dorsal attention A networks in module 3, and the default mode, limbic, and temporoparietal networks in module 4 (*p* < 0.0002).

Using the same approach, we detected two modules in cluster 2 event co-fluctuation patterns. Compared to a spin-test null distribution, module 1 was significantly overrepresented with nodes in the dorsal attention, salience ventral attention, somatomotor, visual networks, and module 2 overrepresented with control, default mode, limbic, and temporoparietal networks (*p* < 0.0002). The modular organization of the average cluster 1 and cluster 2 event co-fluctuation patterns were visualized in Fig. S5, after reorganizing nodes by modules. Also, we find that the modular organization of cluster 2 events largely align with the sensorimotor-association axis found when using principal component analysis (Fig. S8).

In sum, we find that event co-fluctuation patterns are significantly more modular than non-event frames, whose modularities decrease with age. Our results also show that the modular organizations also vary across event co-fluctuation patterns - with four modules detected in cluster 1 events and two modules in cluster 2 events. Given that event frame contribute disproportionately more to static FC than non-events, and that their modularity is significantly greater than both static FC and non-event frames, their reduction in modularity may be driving changes in functional modular organization across the human lifespan which requires further investigation.

### Predicting cognitive measures using event co-fluctuation patterns

To this point, our analyses addressed how we can detect event co-fluctuation patterns and characterized their features across age and coupling with structural connectivity. In this final section, we addressed whether event co-fluctuation patterns are useful in predicting cognitive measures using connectome-based predictive modeling or CPM [57]. Here, we used the Wechsler Individual Achievement Test (WIAT) and Wechsler Abbreviated Scale of Intelligence (WASI) which are various measures of human intelligence or achievement that were available to most subjects across all age groups. Overall, we find that across these different measures, event pattterns consistently yielded stronger predictions of behavioral phenotypes (0.351-0.469), compared to non-event frames (0.335-0.359; Fig. 7). However, we note that the predictions using events or non-events (which uses only 0.1*∼*1% of temporal dynamic information) have weaker correlations than compared to static FC in which these correlations were between 0.66 up to 0.71 (Fig. S6).

**FIG. 7.**
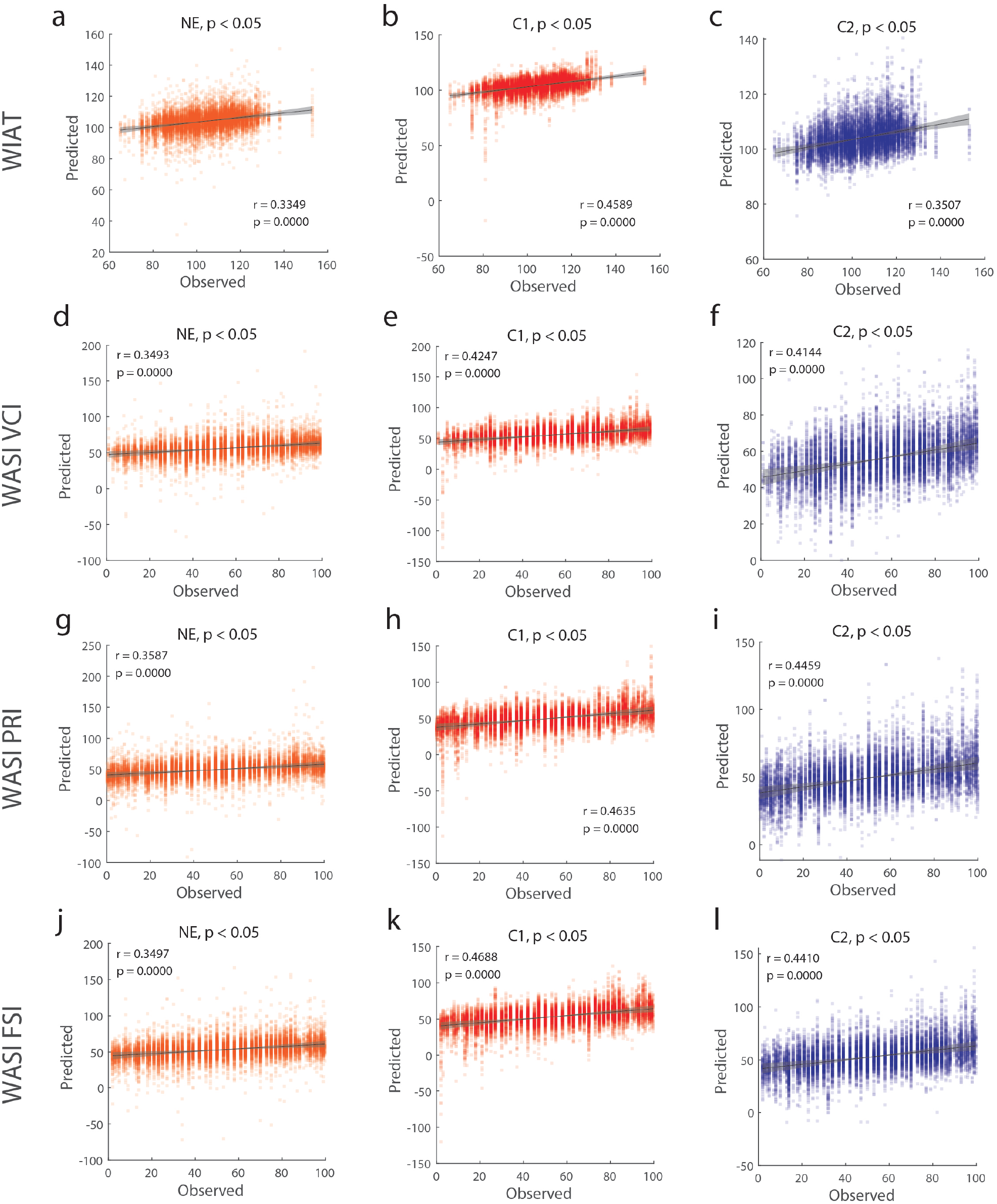
Predicting phenotypes with event and non-event frames using connectome-based predictive modeling (CPM) (*a-c*) Prediction results of WIAT scores using non-event, cluster 1, and cluster 2 frames. (*d-f*) Prediction results of WASI VCI scores using non-event, cluster 1, and cluster 2 frames. (*g-i*) Prediction results of WASI PRI scores using non-event, cluster 1, and cluster 2 frames. (*j-l*) Prediction results of WASI FSI scores using non-event, cluster 1, and cluster 2 frames.

## DISCUSSION

In this study, we aimed to uncover how high-amplitude co-fluctuations – “events” – in resting state functional MRI varied across the human lifespan using a large, cross-sectional sample dataset. First, we demonstrated that events could be partitioned into two clusters based on their topography. We showed that the relative frequency of each cluster varied systematically with age. Next, we aimed to understand the structural underpinnings of these patterns. We addressed this question by using stylized communication models, showing that event co-fluctuations were best explained by geometry (interregional Euclidean distance) and communicability. Moreover, we found that the explanatory power of these predictors varied with age in each pattern. Lastly, we characterized the modular organization of the event co-fluctuation patterns and demonstrated their utility in predicting phenotypes. We showed that event co-fluctuation patterns exhibited dissociable modular organization and that they enhanced the prediction accuracy of scores of WIAT (Wechsler Individual Achievement Test) and WASI (Wechsler Abbreviated Scale of Intelligence) compared to the non-event counterparts.

### Brain-wide events can be clustered into two distinct patterns that change with age

FC is known to continuously undergo refinement across the human life span - from childhood, adolescence, adulthood, and aging [8, 10–12, 19]. Also, previous research has shown that events contribute disproportionately to static FC [24] and can be partitioned into recurring states [45]. In this section, we aimed to answer explore what drives lifespan changes in FC by studying clusters of events.

We also note that patterns of events in part align with and in part vary from previous research. Prior studies that cluster high-amplitude co-fluctuation frames in resting state fMRI report of a task-negative event co-fluctuation pattern that highly correlates with static FC [24, 33, 45], which aligns with the cluster 2 events. In case of the dissimilarities, there may be various reasons underlying the differences in event co-fluctuation patterns.

For one, the event co-fluctuation patterns detected in this investigation were equally sampled across age groups, which may vary from patterns of events found in datasets with scopes limited to healthy young adults [58] or a single densely sampled individual [37]. Also, there were methodological differences in event co-fluctuation pattern detection such as modularity maximization in previous studies [37, 58]. In this study, we used K-means clustering to detect recurrent patterns or states in high amplitude co-fluctuations. This was a methodological decision since our dataset, unlike the previous studies, included over 500 subjects which would have been computationally challenging if we were using modularity maximization.

We also note that previous studies that cluster patterns or states in time-varying, dynamic functional connectivity using sliding window approaches also report varying numbers of recurrent states in resting state fMRI [59–63]. This may be due to the differences in the datasets (e.g. age range), methods to identify states, and MR processing pipelines. Therefore, identifying a consistent set of recurrent states in resting state fMRI requires further investigation. Our results also align with previous findings in dynamic FC which have demonstrated the complexity of dynamic FC decreasing with age by the changes in state dwell times or the slowing of fluctuations [64, 65]. Cluster frequencies as well as their system-level organization were found to vary across age groups. The static FC-resembling, internally similar, S-A axis aligned cluster 2 events showed significant increase in frequency amongst events with age whereas the non-FC-resembling, internally dissimilar, cluster 1 events decreased with age. We can combine the fact that mathematically, events will contribute disproportionately more to static FC [24] and our finding that event co-fluctuation pattern frequencies vary across age groups. Combining these effects, the occurrence of these event co-fluctuation patterns may explain differences in static FC across age groups through a systematic refinement of functional connectivity.

In cluster 1, the system-level changes were largely associated with the reduced co-fluctuation amplitudes in the visual networks across age and increased co-fluctuation amplitudes in somatomotor networks. In cluster 2, there were mainly reduced co-fluctuation amplitudes in the default mode network with age and increases in dorsal attention, limbic, salience ventral attention, somatomotor, temporoparietal, and visual networks with age. These results suggest that events that disproportionately contribute to static FC have distinctive patterns that heterogeneously change with age, effects of which may be diluted when using the entire time series data as in static functional connectivity.

Lastly, we note that it was a decision to focus on these high-amplitude frames since they *by definition* will contribute more to the time-averaged static FC. Recent studies have shown that frames of various amplitudes of co-fluctuation contain different predictive utilities for phenotypes [45, 66–68]. Therefore, we consider zooming in on event frames as a starting point in the analysis of patterns in time-varying co-fluctuations, rather than as a complete summary.

### Structural connectivity derived communication policies and their coupling with event co-fluctuation patterns change with age

Understanding the interplay between structural connections constraining and facilitating synchronized interregional activity, is a central question in neuroscience [47, 69, 70]. To investigate their relationships, we used stylized network models of brain communication to examine the structural underpinnings of event co-fluctuation patterns. We used various communication policies [40] by transforming a sparse SC matrix into fully-weighted matrices. We note that various approaches have previously been used to study SC-FC relationships using neural mass models (NMMs) [71–74] and various heuristics applied to structural connective weights [75–77]. While these approaches have been useful, each branch of methodology is also limited by heavy computation and extensive parameter fitting or being unable to predict FC that aren’t directly connected structurally.

The SC-derived communication measures allowed us to more directly compare structure and function matrices through the use of measures that embody different nuanced policies of communication without the computational burden of multi-parameter models. First, we found that the event co-fluctuation patterns couple with various SC-derived communication measures heterogeneously. We note, that the patterns of explained variance were largely similar across various communication policies. Cluster 1 events showed greatest levels of explained variance in visual areas whereas in cluster 2 events the default mode networks showed greatest levels of explained variance.

These results align with previous research in structure-function coupling in dynamic FC which have demonstrated greater SC-FC coupling in sensorimotor cortices and weaker coupling in the heteromodal regions [39, 78–80]. Such effects may in part be attenuated by the increased frequency of cluster 2 patterns in static FC with increasing age, which were also significantly aligned to the sensorimotor-association axes [81]. However, it is left for further investigation on whether repeated events across the course of the human lifespan affects structure-function relationships.

Next, our results also demonstrated that structural connectivity-derived communication measures and their explained variance for event co-fluctuation patterns change with age. Overall, our results are in line with previous studies that report the relationship of spatial, geometric distance between brain regions [82–84], the topological organization in SC and its relationship with FC [48, 49], and their combinations [85]. Previously, Euclidean distance and communicability was emphasized in describing SC-FC relationships across the human lifespan [40]. In this paper, we demonstrate that both the spatial and topological organization of SC are useful measures for understanding the SC-event relationships across the human lifespan.

Thirdly, we showed that SC-event relationships with age are heterogeneous for event co-fluctuation patterns. A previous study has shown that global SC-FC coupling largely decreases with age and that local SC-FC coupling is heterogeneous across the human lifespan [40]. For both cluster 1 and cluster 2 events, both Euclidean distance and communicability generally showed system-level decreases in coupling with age. Such results align with previous studies in that events disproportionately resemble static FC and therefore are likely to match or even drive the results found in static FC. When investigating the relationship between the event co-fluctuation patterns and Euclidean distance or communicability, the somatomotor networks and visual networks displayed the highest levels of explained variance. However, how these relationships differ across age groups were heterogeneous, and mostly negatively correlated with age. These observations pose further questions for research toward the relationship between structural and functional connectivity and how their relationships are modulated across the human lifespan.

### Event co-fluctuation patterns have distinct modular organizations

The functional organization of the human brain is known to have modular organization, the modularity of which is known to largely decrease with age [52, 86–90]. Here, we extended our study to investigate whether event co-fluctuation patterns are modular and to track their changes in modularity with age. First, we found that as in previous studies using static FC, modularity of event frames were found to decrease with age. Also, the modularity of static FC was found to be driven by that of events, with event co-fluctuation patterns being more modular than static FC and non-event frames. We also found that the event co-fluctuation patterns’ modularity weighed by their relative frequencies of each age group also decrease in modularity across the human lifespan. This result aligns with our expectations since static FC is calculated by averaging over all time frames, including both event and non-event frames.

We also found that each event co-fluctuation pattern has varying modular organizations with different functional brain organizations. The modules in cluster 1 were found to heterogeneously align with canonical functional systems [91] whereas the modules in cluster 2 largely partitioned the brain into higher-order versus lower-order functioning brain regions [81]. Whether the functional organization of the brain is attentuated by the occurrences and organization of high-amplitude co-fluctuations across the human lifespan requires further investigation.

### Event co-fluctuation patterns contain disproportionately more information of an individual than non-events in certain measures

Previous analyses focused on detecting and describing the two main patterns in high-amplitude co-fluctuations in fMRI data. To the best of our knowledge, our analyses are the first to investigate these events and their patterns across the human lifespan. However, our results still beg a practical question: why should one take interest in these events and are they useful for making meaningful predictions of an individual? To address these questions, we used an individual’s average event co-fluctuation pattern as the predictive functional connectome in connectome-based predictive modeling [57] to determine their utility in predicting one’s cognitive performance in achievement and intelligence.

When using cluster 1, cluster 2 event co-fluctuation patterns, and non-events, events were found to outperform predicting measures of achievement and intelligence (the Wechsler Individual Achievement Test and Wechsler Abbreviated Scale of Intelligence scores) compared to non-events. However, we note that when using static FC most clearly outperformed events or non-events, which included at least 100 and up to 1000 times the temporal information as the counterparts. This result is also in line with more recent studies revealing that the high-amplitude co-fluctuations include group-relevant task variance [92] and if removed, can improve subject identifiability [93]. This does not necessarily conflict with our findings that event frames are more identifiable than non-events. When comparing against various co-fluctuation amplitudes, sub-event frames have been reported to be the most identifiable than non-events [66, 68]. Therefore, while event frames are significantly more identifiable than non-event frames, they may also be including group-level variance that does not contribute to subject identifiability.

Additionally, these results hint at the varying utility of event co-fluctuation patterns and dynamic functional connectivity. For predicting achievement and intelligence scores, cluster 1 events produced phenotype predictions that were most similar to the observed phenotypes. However, when determining subject idiosyncrasies, cluster 2 events were found to be more similar within each subject than cluster 1 or non-events. These findings indicate that different event co-fluctuation patterns have varying utility and may even have different functionalities in brain function and throughout the human lifespan. However, we note that investigating event co-fluctuation patterns for predicting phenotypes is a much more limited scope than investigating various moments of co-fluctuation amplitudes in time to maximize predictability [66–68]. How the predictability of various moments of co-fluctuation amplitudes change across the human lifespan requires further investigation.

### Limitations and future directions

Finally, we highlight some of the limitations in our present study. First, our results mainly describe event co-fluctuation patterns and their changes with age but does not provide an understanding of their roles in our brains’ activities. Our results revealed that events have distinct subtypes that change throughout the human lifespan, are heteromodally involved with SC-derived communication measures, are modular, and have predictive utility of one’s achievement and intelligence. However, it still remains unclear whether an event co-fluctuation pattern serves specific functional roles such as development, functional diversification, stabilizing and/or sustaining the main functions of the human brain. Also, it is unknown whether event co-fluctuation patterns are involved in the manifestation of cognitive or behavioral disorders through their changes. Further investigation is required to determine the relationship between each event co-fluctuation pattern with brain function and age. Another question that has not been resolved in this paper is what causes or drives these patterns. Previous studies have shown that high-amplitude co-fluctuations in fMRI are related to endogenous hormones during a menstrual cycle [37], may be implicated in arousal during movie watching data [94], can be predicted by static FC [31], and arise in modular in silico activation patterns [41]. In part, we aimed to answer this question by investigating the relationship between event co-fluctuation patterns and various SC-derived communication measures. However, a longitudinal investigation is warranted to help understand the structure-function relationships that supports the event co-fluctuation patterns.

Another limitation of our study is that we mainly chose to describe lifespan trajectories of variables of interest with linear models using linear correlation coefficients. While linear models have been widely used to observe age-related changes in brain networks, they have been known to be prone to some inherent limitations [95, 96]. One which includes that first order polynomial models may not capture nonlinear changes in the variables over the lifespan. In future work, other models such as partial least squares may prove useful for identifying collective changes in connectivity with age [97].

Also, the number of clusters identified in this study requires further investigation since this may vary depending on the method of cluster detection and error estimation, dataset, or even due to variance in the data preprocessing pipelines. In this study, our decision to use the K-means algorithm was practically motivated since methods used in previous studies with smaller datasets was computationally challenging to scale to the NKI dataset including hundreds of subjects.

Lastly, the dataset that we investigate covers a limited age range (ages 6 to 75), which does not cover the early postnatal-childhood period (ages 0 to 5) nor healthy aging subjects beyond the age of 75. It can be expected that the young brains of infants and children also experience such high-amplitude co-fluctuations in some form, and their occurrences may be related to the observed events in their later years. Also, it is likely that due to the rapid change in structural and functional connectivity in the early years, that such event co-fluctuation patterns also undergo dynamic changes during this period.

## Conclusions

In conclusion, our work sought to answer whether high-amplitude co-fluctuations in the human BOLD signal has patterns that consistently show up across age which are also subject to change across the human lifespan. Our findings show that events or peak moments of high-amplitude co-fluctuations in rsfMRI, which constitute 0.1*∼*1% of global time series, have distinct patterns that change throughout the human lifespan. These patterns were also found to change in their coupling patterns with one’s structural connectivity-derived communication measures and to have distinct modular organizations. Finally, we demonstrate that event frames are more predictive of an individual’s achievement and intelligence scores across age than non-event frames, highlighting their potential predictive utilities in future research.

## MATERIALS AND METHODS

### Dataset

#### Nathan Kline Institute, Rockland Sample

The Nathan Kline Institute Rockland Sample (NKI-RS) dataset consisted of resting state functional magnetic resonance imaging, structural magnetic resonance imaging, as well as diffusion magnetic resonance imaging data from 711 subjects (downloaded December 2016 from the INDI S3 Bucket) of a community sample of participants across the human lifespan. After excluding subjects based on data and metadata completeness and quality control (see Image Quality Control), the final subset used included 537 subjects (62.6% female, age range = 6 - 75). The study was approved by the Nathan Kline Institute Institutional Review Board and Monclair State University Institutional Review Board and informed consent was obtained from all subjects. Subjects were compensated for their participation. A comprehensive description of the imaging parameters can be found online at the NKI website. Briefly, images were collected on a Siemens Magneton Trio with a 12-channel head coil. Subjects underwent three differently parameterized resting state scans, but only one acquisition is used in the present study. The fMRI data was acquired with a gradient-echo planar imaging sequence (TR = 645ms, TE = 30ms, flip angle = 60^*◦*^, 3mm isotropic voxel resolution, multiband factor = 4). This resting state run lasted approximately 9:41 seconds, with eyes open and instructions to fixate on a cross. Subjects underwent one T1-weighted structural scan (TR = 1900ms, TE = 2.52 ms, flip angle = 9^*◦*^, 1mm isotropic voxel resolution) and one diffusion MRI scan (TR = 2400ms, TE = 85ms, flip angle = 90^*◦*^, 2mm isotropic voxel resolution, 128 diffusion weighted volumes, b-value = 1500s/mm^2^, 9 b = 0 volumes).

### Image Quality Control

The NKI was downloaded in December of 2016 from the INDI S3 Bucket. At the time of download, the dataset consisted of 718 fMRI (“acquisition645”; 634 subjects) 957 T1w (811 subjects), and 914 DWI (771 subjects) images. fMRI images were excluded if greater than 15% of time frames exceeded 0.5mm framewise displacement. Furthermore, fMRI images were excluded if the scan was marked as an outlier (1.5x the inter-quartile range in the adverse direction) in 3 or more of the following quality metric distributions: DVARS standard deviation, DVARS voxel-wise standard deviation, temporal signal-to-noise ratio, framewise displacement mean, AFNI’s outlier ratio, and AFNI’s quality index. This image quality metric filtering excluded 21 fMRI images, zero T1w images, and 16 DWI images. Following these visual and image quality metric filterings, 697 fMRI images (633 subjects), 809 T1w images (699 subjects), and 728 DWI images (619 subjects) were maintained.

### Image Preprocessing

The fMRI images in the NKI dataset were preprocessed using the fMRIPrep version 1.1.8 [98]. The following description of fMRI preprocessing is based on fMRIPrep’s documentation. This workflow utilizes ANTs (2.1.0), FSL (5.0.9), AFNI (16.2.07), FreeSurfer (6.0.1), nipype [99], and nilearn [100]. T1w images were submitted to FreeSurfer’s cortical reconstruction workflow (version 6.0). The FreeSurfer results were used to skull strip the T1w, which was subsequently aligned to MNI space with 6 degrees of freedom. Functional data was slice time corrected using AFNI’s `3dTshift` and motion corrected using FSL’s `mcflirt`. “Fieldmap-less” distortion was performed by co-registering the functional image to the same-subject T1w with intensity inverted [101] constrained with an average fieldmap template [102], implemented with `antsRegistration`. This was followed by co-registration to the corresponding T1w using boundary-based registration [103] with 9 degrees of freedom, using `bbregister`. Motion correcting transformation, field distortion correcting warp, and BOLD-to-T1w transformation warp were concatenated and applied in a single step using `antsApplyTransforms` using Lanczos interpolation. Frame-wise displacement [104] was calculated for each functional run using Nipype. The first four frames of the BOLD data in the T1w space were discarded. Each T1w was corrected using `N4BiasFieldCorrection` [105] and skull-stripped using `antsBrainExtraction.sh` (using the OASIS template). The ANTs derived brain mask was refined with a custom variation of the method to reconcile ANTs-derived and FreeSurfer-derived segmentations of the cortical gray matter of Mindboggle [106]. Brain tissue segmentation of cerebrospinal fluid (CSF), white matter (WM) and gray matter(GM) was performed on the brain-extracted T1w using `fast` [107]. Diffusion images were preprocessed following the “DESIGNER” pipeline using MRTrix (3.0) [108, 109], which includes denoising, Gibbs ringing and Rician bias correction, distortion and eddy current correction [110] and B1 field correction. DWI were then aligned to their corresponding T1w and the MNI space in one interpolation step with B-vectors rotated accordingly. Local models of white matter orientation were estimated in a recursive manner [111] using constrained spherical deconvolution [112] with a spherical harmonics order of 8. Tractography was performed using Dipy’s `Local Tracking` module [113]. Probabilistic streamline traactography was seeded five times in each white matter voxel. Streamlines were propagated with a 0.5mm step size and a maximum turning angle set to 20^*◦*^. Streamlines were retained if longer than 10mm and with valid endpoints, following Dipy’s implementation of anatomically contrained tractography [114].

### Network definition

#### Parcellation

For the NKI fMRI and DWI, the Schaefer 400 parcellation was rendered as a volumentric parcellation in each subject’s anatomical space within the gray matter ribbon. To transfer the parcellation from fsaverage to subject space, FreeSurfer’s `mris ca label` function was used in conjunction with a pre-trained Gaussian classifier surface atlas [115] to register cortical surfaces based on individual curvature and sulcal patterns.

#### Functional connectivity

For the NKI dataset, each preprocessed BOLD image was linearly detrended, band-pass filtered (0.008-0.08Hz), confound regressed, and standardized using Nilearn’s `signal.clean` function, which removes confounds orthogonally to the temporal filters. The confound regression strategy included six motion estimates, mean signal from the white matter, cerebrospinal fluid, and whole brain mask, derivatives of these previous nine regressors, and squares of these 18 terms. Spike regressors for frames with motion greater than 0.5mm framewise displacement were applied. The 36 parameter strategy (with and without spike regression) has been shown to be a relatively effective option to reduce motion-related artifacts [116]. Following these preprocessing operations, the mean signal was acquired for each node in the volumetric anatomical space.

#### Structural connectivity

Structural connectivity was quantified based on the number of streamlines between cortical regions (nodes). Since the size of the node is has known effect on the streamline count [75], the streamline counts were normalized by dividing the count between nodes by the geometric average volume of the nodes.

### Age matched sampling and binning

The intersection of subjects with at least one valid fMRI, T1w, and DWI images after totaled in 567 subjects. Age metadata was available for 542 of these subjects. Finally, subjects with fMRI images with NAN values were excluded, resulting in an intersection of 537 subjects (age 6 - 75). The age distribution of the NKI dataset was not uniformly distributed, which if used directly for the clustering analysis may bias the cluster results to characteristics of age groups with larger samples. Therefore, we first created seven equal sized age bins (bin size = 10 years) and randomly sampled 20 subjects per age group. The number of subjects randomly sampled in each age group was determined to ensure that the age group with the smallest number of N could be sampled to include on average less than a 50% overlap in any pair of random samples. Each subject’s nodal time series were used to calculate edge time series and detect events. The process of subject sampling, event detection, and clustering processes were repeated 100 times.

### Edge time series and events

Following the preprocessing and sampling steps for rsfMRI data described previously, the mean signal was taken at each time frame for each node, forming the nodal time series. The FC between brain regions *i* and *j* is operationalized as a correlation coefficient summarized as the Pearson correlation coefficient as follows:

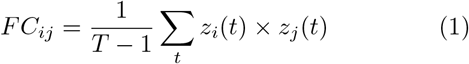

where *z*_*i*_ = [*z*_*i*_(1), …, *z*_*i*_(*T*)] is the vector or z-scored nodal activity from region *i*.

The edge time series for edge *i, j* is calculated by simply omitting the summation and normalization step. In short, edge time series is calculated as follows:

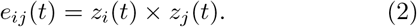

This procedure is repeated for all pairs of nodes resulting in an edge-by-time time series matrix. The elements of this matrix encode the moment-by-moment co-fluctuation magnitude of nodes *i* and *j*. A positive value in this co-fluctuation would indicate a simultaneous increase or decrease in the activity of nodes *i* and *j*, whereas a negative value would reflect their opposite direction of activity. Similarly, a magnitude close to zeros would indicate that either *i* or *j* had very low levels of activity.

After creating an edge time series matrix (edge-by-time) for each subject, we calculated the root mean sum square (RMS) at every given time point resulting in a single time series representing the global co-fluctuation amplitude. Next, we identified frames as “events” in the RMS signal that had a significantly larger RMS than the circularly shifted null model counterpart.

### K-means clustering

We used a k-means clustering algorithm with both Pearson correlation and Lin’s concordance as the distance measure to cluster the event co-fluctuation patterns. More specifically, events frames were partitioned in a non-overlapping fashion so that each frame was labeled either as cluster 1 up to cluster K. We acquired the event frame clusters for each run from k = 2 to 10. The partition labels in k-means are assigned randomly - the identical set of elements can have an identical partition with alternative labels [*C*1, *C*1, *C*1, *C*2, *C*2, *C*1] or [*C*2, *C*2, *C*2, *C*1, *C*1, *C*2]. Therefore, we realigned the cluster labels across runs so that each cluster label represented the maximally similar cluster label in another run. To do align the cluster labels, at each K, the cluster labels were compared across runs and realigned to minimize a cost function. We used the *matchpairs* function provided in MATLAB that minimizes total cost - measured as cluster centroid dissimilarity (1 − Correlation coefficient) - of a linear assignment problem. Cluster centroids from each run were then re-aligned to match the centroids of the partition that minimizes the total cost.

### SC predictors

A suite of communication measures (predictors) were applied to each subject’s structural connectivity matrices to help uncover the SC-event co-fluctuation pattern relationships. A total of 34 predictors (a core set of 10 measures with varying weights in communication policy) were used. A more detailed description into all 10 measures can be found in Zamani Esfahlani et al. 2022 [40].

We focused on two communication measures that were mainly highlighted in our results - Euclidean distance and communicability. Euclidean distance was calculated between regional centers of mass as the square root of squared difference between center coordinates.

Communicability [56] is the weighted sum of all walks between pairs of nodes. For a binary network, communicability is calculated as 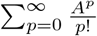. Walks and their weights are dependent on the number of steps, and longer walks are penalized in their contributions. For instance, a single step walks are 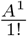, two-step walks are 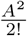, three-step walks are 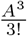, and so on.

For weighted networks, we calculated weighted communicability by following Crofts and Higham (2009) [117]. First, the weighted structural connectivity matrix is normalized as *A*^*′*^ = *D*^−1*/*2^*AD*^−1*/*2^ (*D* : degree diagonal matrix). The normalized matrix is the exponent used to calculate weighted communicability *G*_*wei*_ *= e*^*A*^*’*.

### Event frame modularity

We used modularity maximization to find modular structures and calculate the modularity of event co-fluctuation patterns, non-event frames, and static FC. Modularity maximization is a computational heuristic for detecting community structures in a network. The method defines communities or clusters as groups of elements in which the internal density of connections maximally exceed what would be expected. Based on this approach, we used the expected weight of connections to be equal to the mean similarity between all pairs of patterns. Here, we used modularity maximization with the generalized Louvain heuristics which is non-deterministic, and can yield dissimilar results depending on the initial state. Therefore, after detecting modules of event co-fluctuation patterns, the partitions were realigned to the cluster centroid that minimizes the dissimilarity cost function after averaging the event co-fluctuation pattern for each age group. We then calculated the co-assignment probability of nodes, i.e. the likelihood that the nodes are assigned to the same community. We repeated the algorithm for 1000 iterations with varying random seeds. The variability across the iterations were resolved by using a consensus clustering algorithm in which we iteratively cluster the module co-assignment matrix until convergence. The resulting consensus partition assigned each brain region in non-overlapping clusters. The modules of non-event frames were detected with an identical approach.

### Frame-wise identifiability

We calculate *differential identifiability* [93] by using event frames across subjects and first creating a frame-to-frame similarity matrix. Similarity between frames were calculated using correlation coefficients, followed by subtracting the mean within-subject frame similarities minus mean between-subject similarities. Specifically, differential identifiability (*I*_*diff*_) was calculated as *Idiff* = (*Iself* − *Iothers*) *∗* 100.

## Author Contributions

YJ and RFB conceived of study, carried out all analyses, and generated figures. JT carried out analyses and contributed to the project *via* discussion. CS contributed in analysis and discussion. JF processed all imaging data and in discussion. All authors helped revise and write the submitted manuscript.

## Data Availability

All imaging data come from publicly-available, open-access repositories. Human connectome project data can be accessed *via* https://db.humanconnectome.org/app/template/Login.vm after signing a data use agreement. Midnight scan club data can be accessed *via* OpenfMRI at https://openfmri.org/dataset/ds000224/.

## Code Availability

All processing and analysis code is available upon reasonable request.

---

**FIG. S1.**
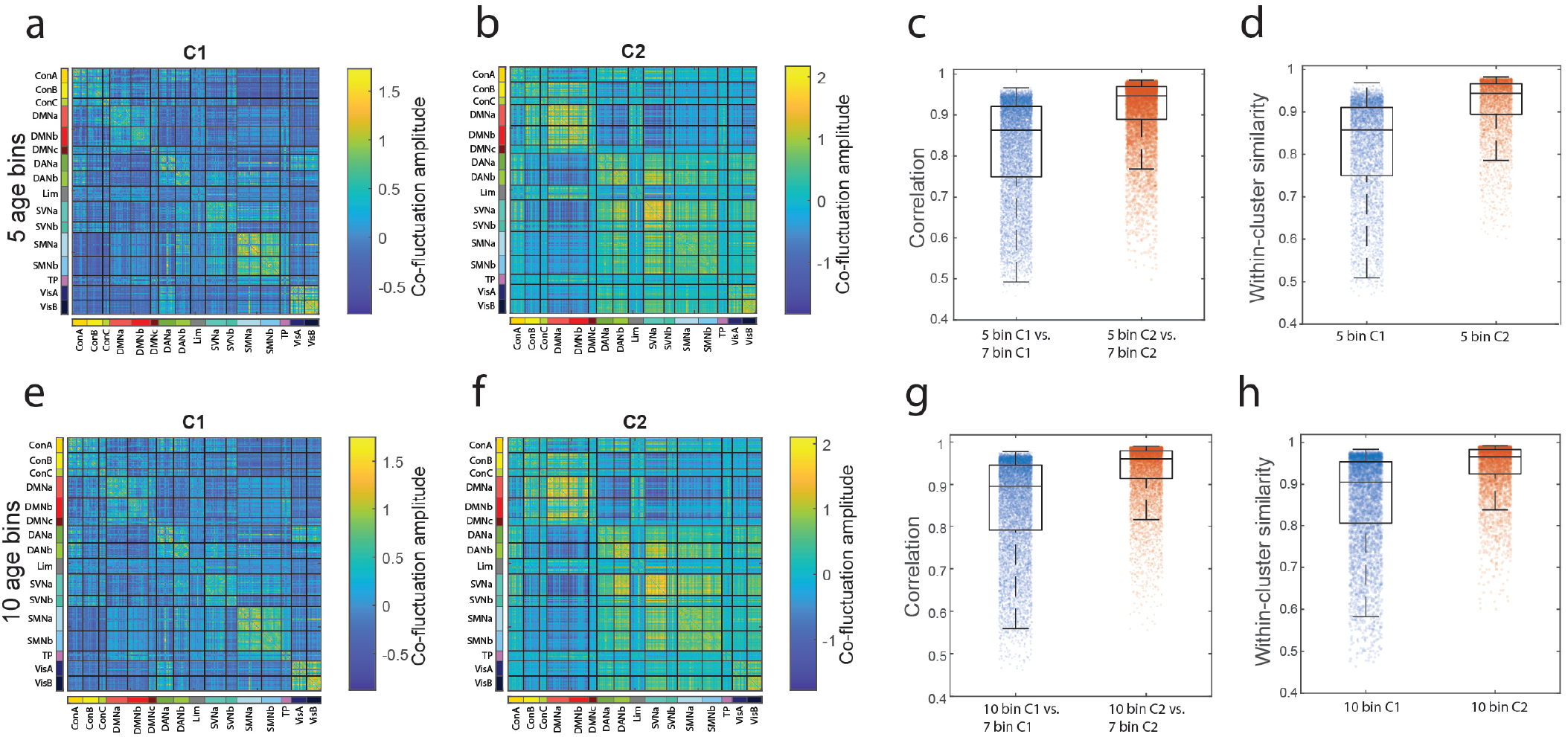
Event co-fluctuation patterns using k-means clustering with alternative age bins. Event co-fluctuation patterns across five age bins (age bin = 14 years). (*a*) Mean C1 and (*b*) mean C2 across all age bins. Relative frequency of (*c*) C1 and (*d*) C2 across all five age bins. (*e*) Similarity of C1 and C2 with static FC for five age bins. Event co-fluctuation patterns across ten age bins (age bin = 7 years). (*f*) Mean C1 and (*g*) mean C2 across all ten age bins. Relative frequency of (*h*) C1 and (*i*) C2 across all ten age bins. (*j*) Similarity of C1 and C2 with static FC for ten age bins.

**FIG. S2.**
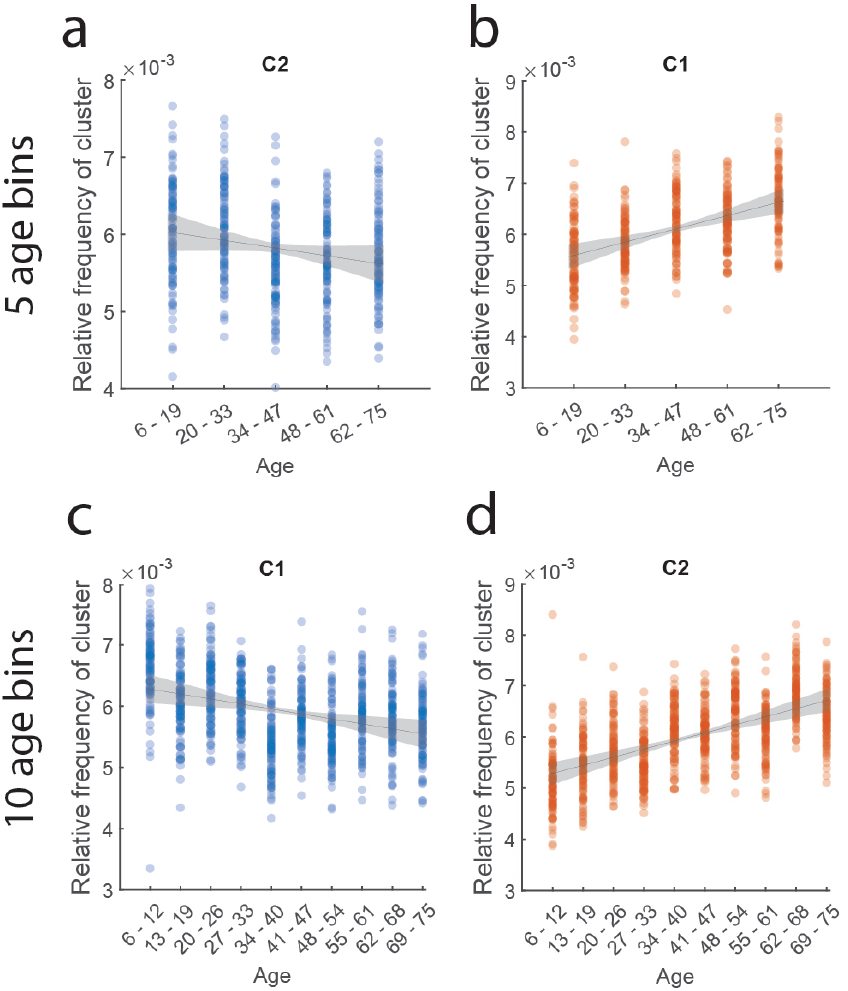
Event co-fluctuation pattern frequencies with age in alternative age bins. Relative frequency of (*a*) C1 and (*b*) C2 across all five age bins. Relative frequency of (*c*) C1 and (*d*) C2 across all ten age bins. (*j*) Similarity of C1 and C2 with static FC for ten age bins.

**FIG. S3.**
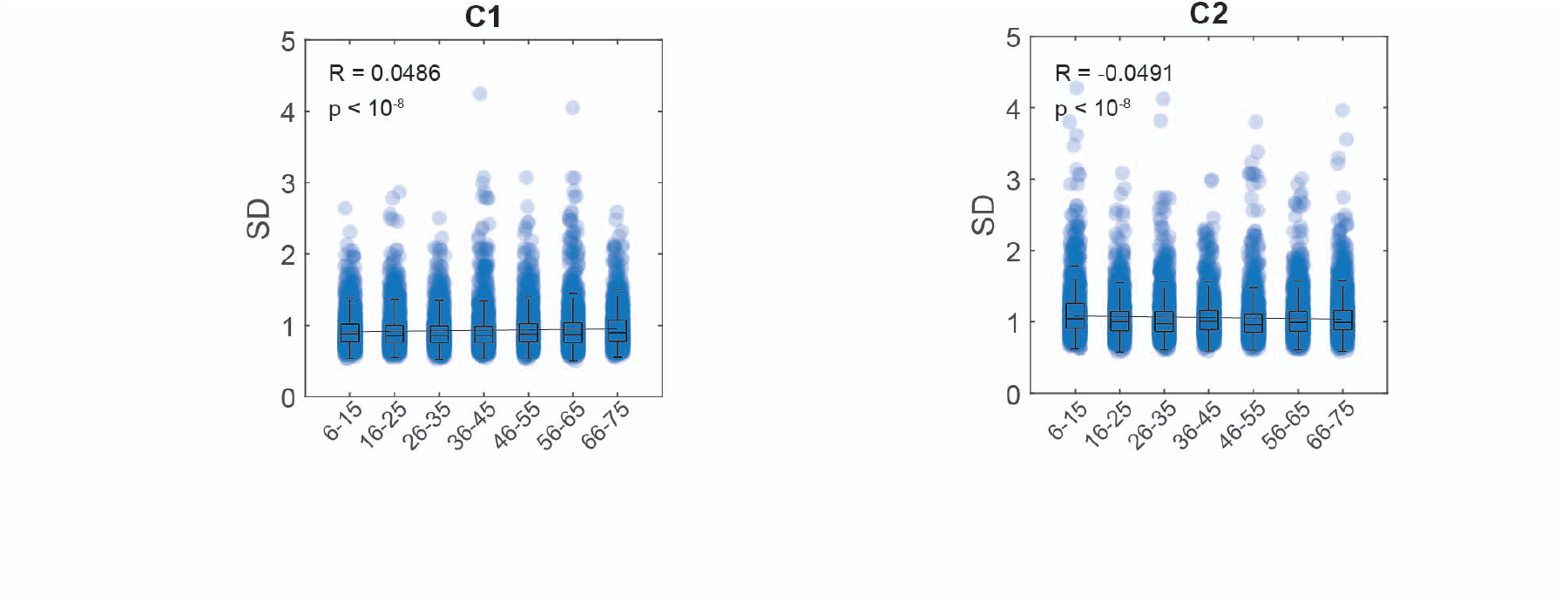
Standard deviation of event co-fluctuation patterns within each age bin. (a) Cluster 1 and their standard deviations across subject events within each age bin across 100 iterations. Correlation of standard deviations of cluster 1 with age (*R* = 0.0486, *p* < 10^−8^). (b) Cluster 2 and their standard deviations across subject events within each age bin across 100 iterations. Correlation of standard deviations of cluster 2 with age (*R* = −0.0491, *p* < 10^−8^).

**FIG. S4.**
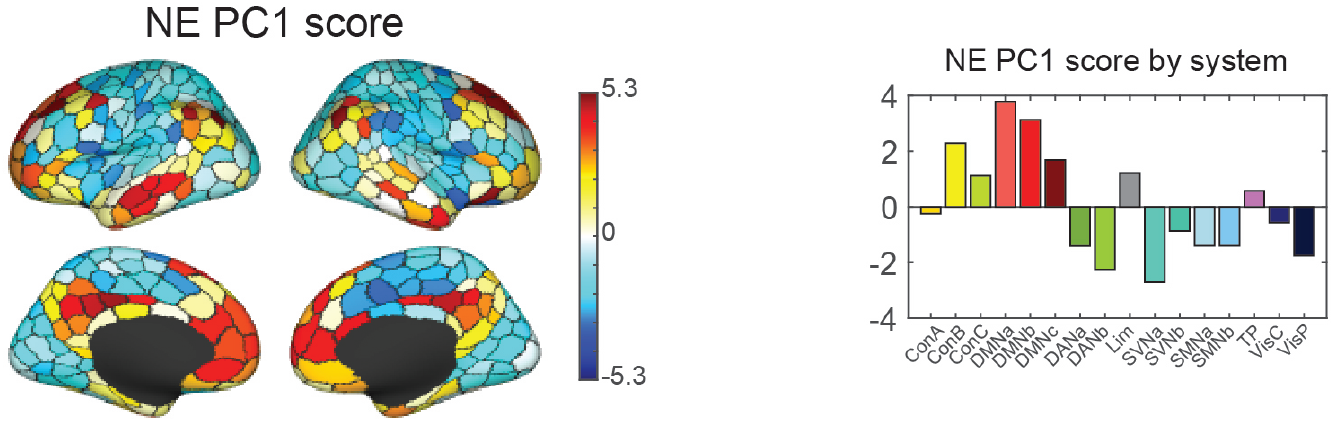
Non-event frames and their first principal component. Brain surface plot and the system-level organization of the first principal component score of the mean non-event frames across age groups.

**FIG. S5.**
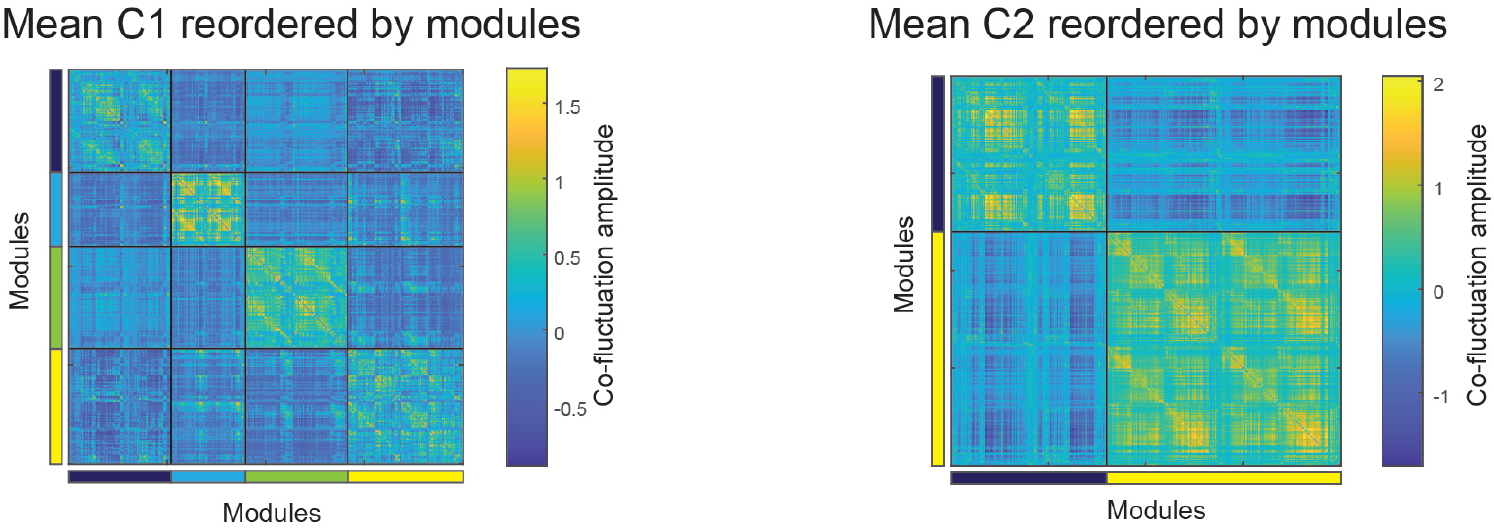
Mean event co-fluctuation patterns reordered by modules. The mean C1 and C2 event co-fluctuation patterns across age groups after reordering nodes by their modular labels.

**FIG. S6.**
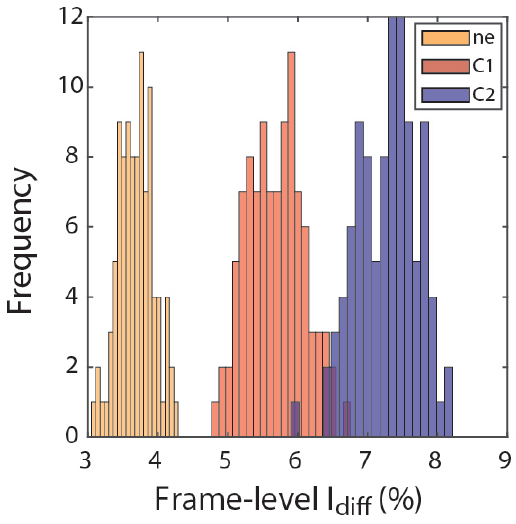
Event co-fluctuation patterns and identifiability. *Differential identifiability* of events (C1, C2) and non-events (NE). Frame-level Idiff calculated as the difference between within-subject cluster frame similarity and between subject cluster frame similarity.

**FIG. S7.**
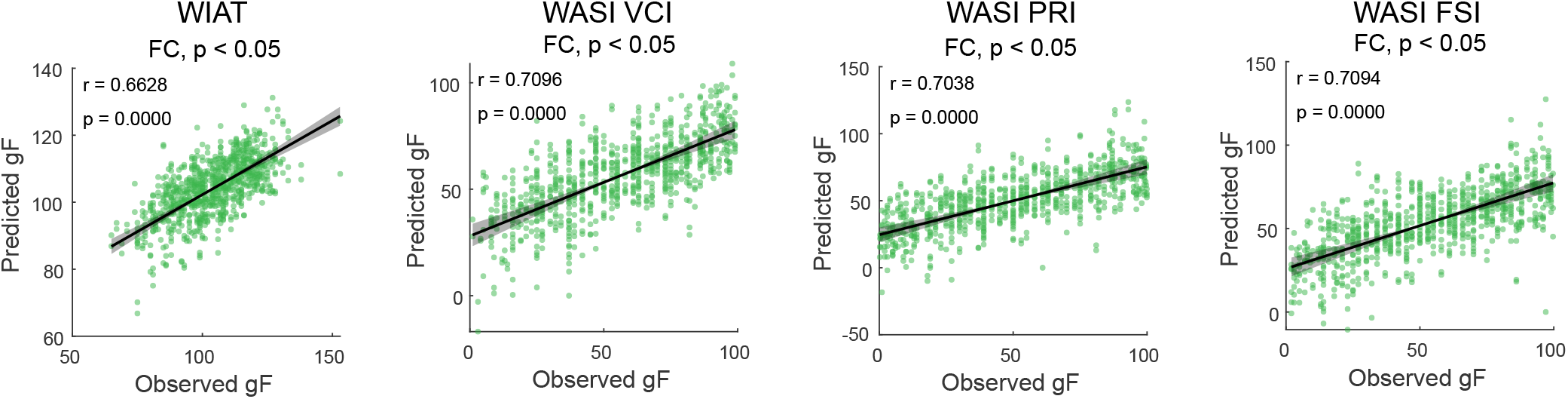
Static FC and predictability of achievement and intelligence scores using CPM. Prediction of WIAT, WASI VCI, WASI PRI, WASI FSI scores based on static FC of individuals using connectome-based predictive modeling (CPM).

**FIG. S8.**
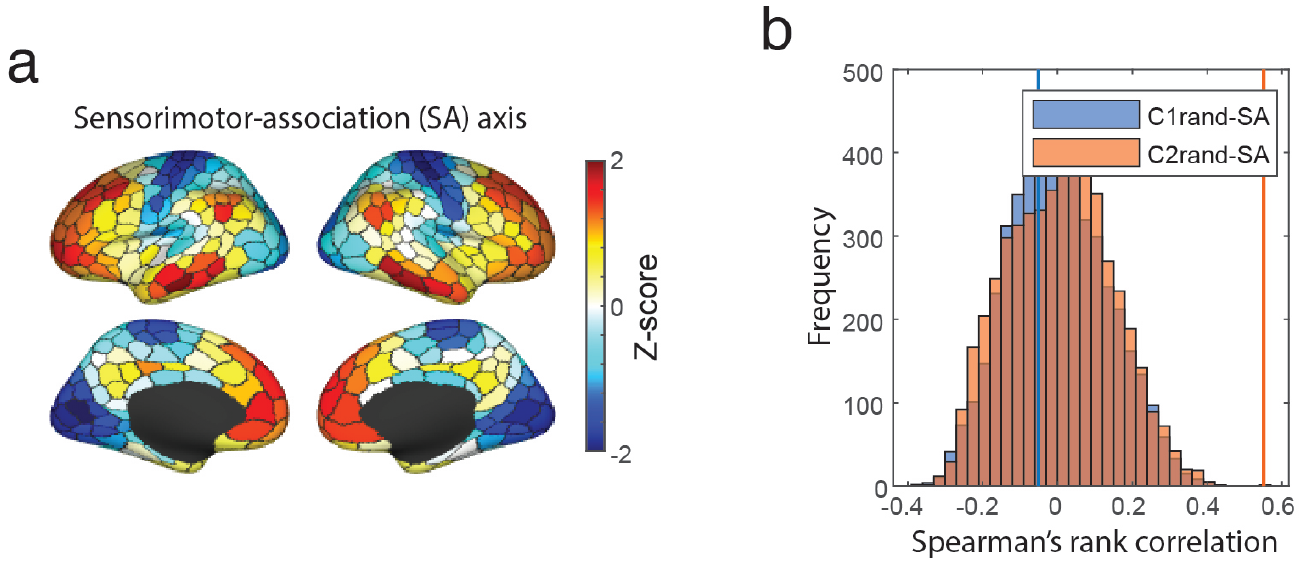
Event co-fluctuation patterns and the sensorimotor-association axis. (*a*) Z-score of the whole brain ranking of nodes on the sensorimotor-association (SA) axis. (*b*) Spearman’s rank correlation between the average event co-fluctuation patterns’ first principal component (PC) and the z-scored global SA axis (C1: orange line; C2: yellow line). Histograms represent the Spearman’s rho between average event co-fluctuation patterns’ first PCs after spin-test nodal randomization (5000 iterations) and the z-scored global SA axis.

**FIG. S9.**
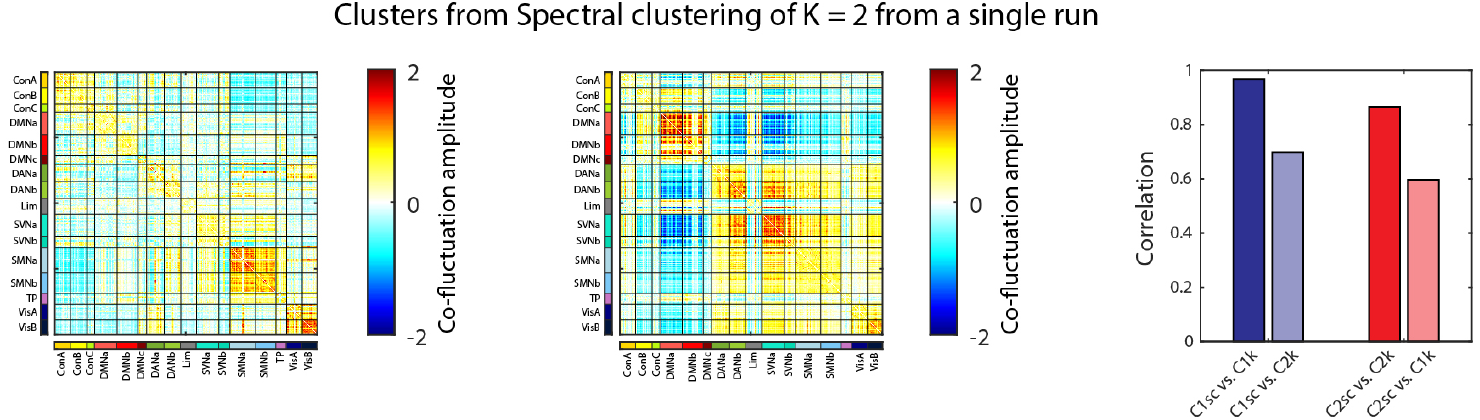
Event co-fluctuation patterns from spectral clustering analysis at K = 2, based on a single sampling process. Event co-fluctuation patterns found using spectral clustering analysis at K = 2 from a single sampling process with cluster 1 (left), cluster 2 (middle), and their correlation with clusters found using K-means clustering (right).

**FIG. S10.**
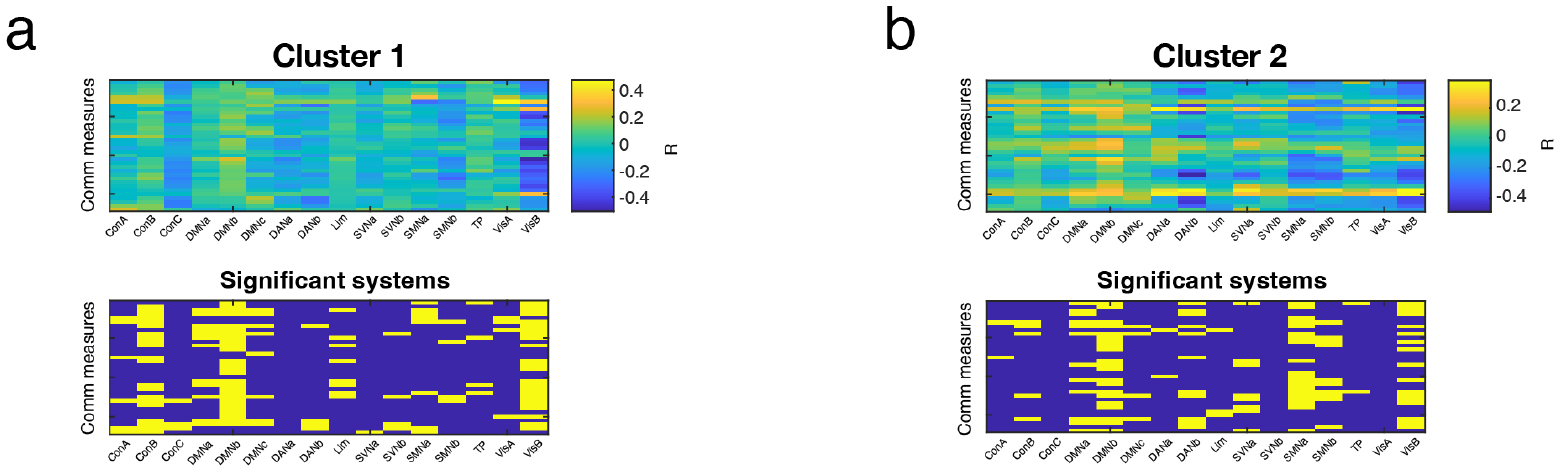
Cluster-SC communication measures relationships with age using age group averages. (*a*) Age correlations of communication measures and systems using age group average estimates of cluster 1 and structural connectivity. (*b*) Age correlations of communication measures and systems using age group average estimates of cluster 2 and structural connectivity.

**FIG. S11.**
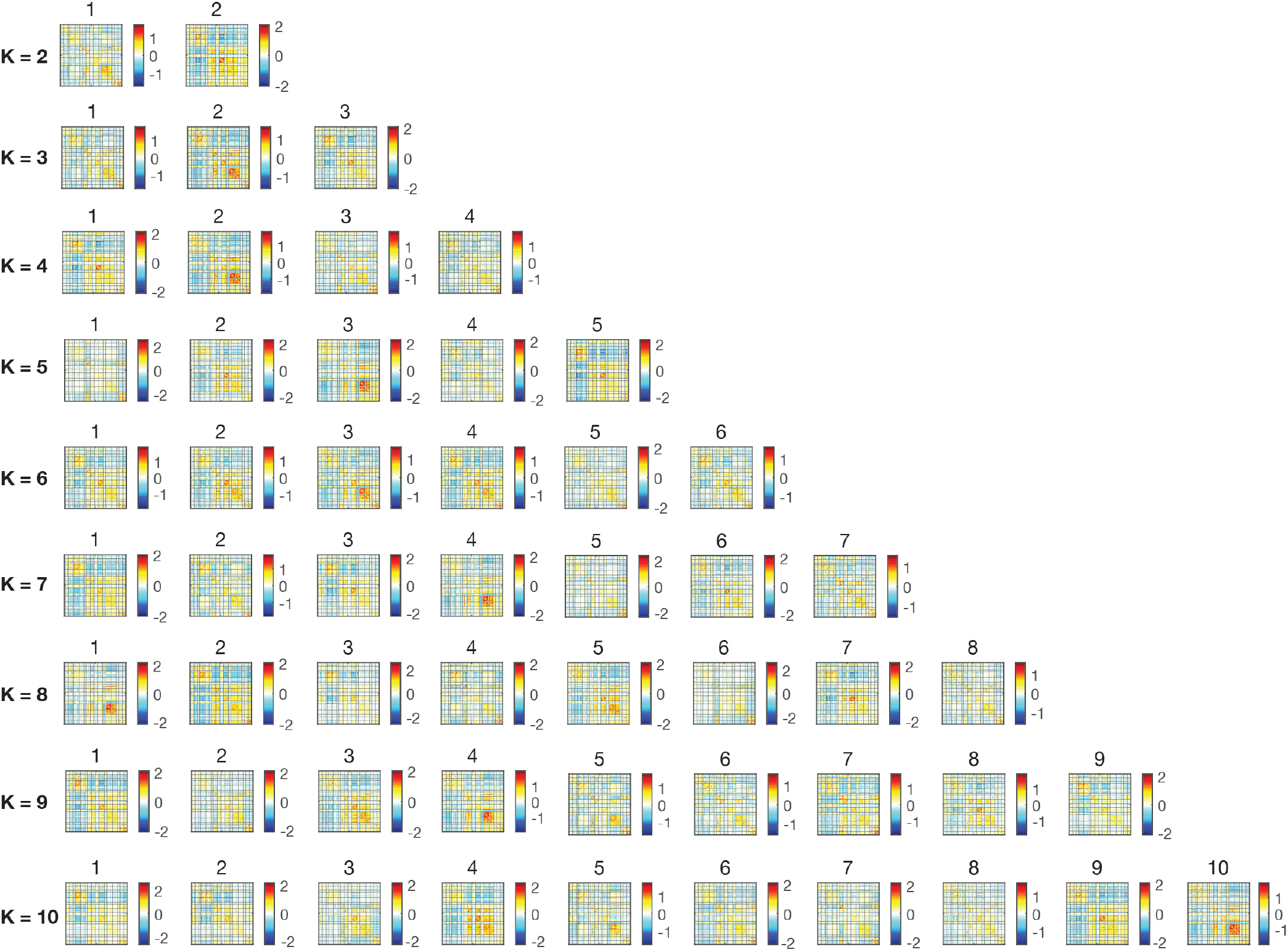
Event co-fluctuation patterns after aligning to cluster centroids at *K* = 2 − 10.

